# SARS-CoV-2 infection damages airway motile cilia and impairs mucociliary clearance

**DOI:** 10.1101/2020.10.06.328369

**Authors:** Rémy Robinot, Mathieu Hubert, Guilherme Dias de Melo, Françoise Lazarini, Timothée Bruel, Nikaïa Smith, Sylvain Levallois, Florence Larrous, Julien Fernandes, Stacy Gellenoncourt, Stéphane Rigaud, Olivier Gorgette, Catherine Thouvenot, Céline Trébeau, Adeline Mallet, Guillaume Duménil, Samy Gobaa, Raphaël Etournay, Pierre-Marie Lledo, Marc Lecuit, Hervé Bourhy, Darragh Duffy, Vincent Michel, Olivier Schwartz, Lisa A. Chakrabarti

## Abstract

Understanding how SARS-CoV-2 spreads within the respiratory tract is important to define the parameters controlling the severity of COVID-19. We examined the functional and structural consequences of SARS-CoV-2 infection in a reconstituted human bronchial epithelium model. SARS-CoV-2 replication caused a transient decrease in epithelial barrier function and disruption of tight junctions, though viral particle crossing remained limited. Rather, SARS-CoV-2 replication led to a rapid loss of the ciliary layer, characterized at the ultrastructural level by axoneme loss and misorientation of remaining basal bodies. The motile cilia function was compromised, as measured in a mucociliary clearance assay. Epithelial defense mechanisms, including basal cell mobilization and interferon-lambda induction, ramped up only after the initiation of cilia damage. Analysis of SARS-CoV-2 infection in Syrian hamsters further demonstrated the loss of motile cilia *in vivo*. This study identifies cilia damage as a pathogenic mechanism that could facilitate SARS-CoV-2 spread to the deeper lung parenchyma.

## INTRODUCTION

The COVID-19 pandemic remains a worldwide public health emergency. The infection caused by the severe acute respiratory syndrome coronavirus 2 (SARS-CoV-2) emerged in Wuhan, China, in December 2019 (Lu et al., 2020; Zhou et al., 2020b; Zhu et al., 2020b) and spread to all continents except Antartica, affecting over 31 million persons and causing over 970,000 deaths as of Sept. 24, 2020 (covid19.who.int).

The respiratory syndrome COVID-19 ranges from mild upper respiratory tract infection to bilateral pneumonia with acute respiratory distress syndrome (ARDS) and multiple organ failure (Guan et al., 2020; Huang et al., 2020; Zhou et al., 2020a). Pathological examination indicates that SARS-CoV-2 targets primarily the airways and the lungs (Tian et al., 2020; Xu et al., 2020). Severe cases are characterized by diffuse alveolar damage and formation of hyaline membranes that limit gaseous exchanges. COVID-19 pneumonia is associated with inflammatory infiltrates in the alveolar space and a systemic cytokine storm, suggesting that an exacerbated immune response contributes to damaged lung function (Huang et al., 2020; Liao et al., 2020). Induction of interferons appears limited in the more severe clinical cases, pointing to an imbalance between antiviral and inflammatory cytokine responses (Blanco-Melo et al., 2020; Hadjadj et al., 2020).

SARS-CoV-2 is genetically closely related to the original SARS-CoV-1 coronavirus, which caused an outbreak of severe acute respiratory syndrome in 2002-2003, affecting about 8,000 patients and resulting in 774 reported deaths (Peiris et al., 2003; Petersen et al., 2020). Both viruses use the angiotensin converting enzyme 2 (ACE2) as a receptor, and the transmembrane serine protease 2 (TMPRSS2) as a viral entry cofactor (Hoffmann et al., 2020; Zhou et al., 2020b). The pneumonia induced by the two coronaviruses show similar features in severe cases. SARS-CoV-2 induces a mortality rate approximately ten times lower than SARS-CoV-1, but shows much higher effective transmissibility, and thus represents a greater threat to global health (Petersen et al., 2020). Possible reasons for the high transmissibility of SARS-CoV-2 include an active viral replication in upper airway epithelia at an early stage of infection. The number of SARS-CoV-2 genomic copies in nasopharyngeal swabs is generally high at symptom onset (≥10^6^ viral RNA copies/mL) and persists for about 5 days before declining (Wolfel et al., 2020; Zou et al., 2020). This is in contrast to the pattern observed for SARS-CoV-1, with viral RNA peaking about 10 days after symptom onset and remaining of moderate magnitude (Peiris et al., 2003). Therefore, analyzing how SARS-CoV-2 spreads in the airways is relevant to understand its pandemic potential and potentially identify novel mitigation strategies.

The epithelium lining the airways plays a key role in the defense against infections (Tilley et al., 2015). It comprises goblet cells that secrete a protective mucus able to trap inhaled particles, including microbes. Ciliated cells, which constitute over half of epithelial cells, possess an apical layer of about 200 cilia that beat rhythmically in a coordinated fashion, resulting in a movement of the overlaying mucus layer towards the laryngopharynx, where it is ultimately swallowed (Spassky and Meunier, 2017). This mechanism of mucociliary clearance prevents the accumulation of particles and mucus within the lungs. The airways basal cells, located close to the epithelial basement membrane, respond to injury by proliferating and differentiating into other epithelial cell types. Studies of autopsy samples from COVID-19 patients and experimental infection of tissue explants have documented SARS-CoV-2 replication predominantly in the upper and lower airway epithelia and in the lung alveoli (Hou et al., 2020; Hui et al., 2020; Yao et al., 2020). Infection of reconstituted airway epithelia showed a preferential targeting of ciliated cells, with an infection of goblet cells in some (Mulay et al., 2020; Pizzorno et al., 2020; Ravindra et al., 2020; Zhu et al., 2020a) but not all studies (Hou et al., 2020; Zhu et al., 2020b). The functional consequences of SARS-CoV-2 infection on epithelial functions remain to be characterized.

To better understand the mechanism of SARS-CoV-2 dissemination in the respiratory tract, we analyzed the ultrastructural and functional changes induced by infection in a reconstituted human bronchial epithelium model. This system enabled the study of SARS-CoV-2 interactions with its primary target cells in a well-differentiated pseudostratified epithelium. We also examined the impact of SARS-CoV-2 infection on the airway mucosa *in vivo*, using the physiologically relevant Syrian hamster model.

## RESULTS

### SARS-CoV-2 actively replicates in a reconstituted human bronchial epithelium model

SARS-CoV-2 infection was studied in the MucilAir™ model, consisting of primary human bronchial epithelial cells grown over a porous membrane and differentiated at the air/liquid interface (ALI) for over 4 weeks (Fig. 1A). We first verified by scanning electron microscopy (SEM) that the bronchial epithelial cells were differentiated into a pseudostratified epithelium comprising three main cell types (Fig. 1B): ciliated cells harboring a dense layer of 5-10 μm long motile cilia, goblet cells enriched in mucus-containing vesicles (arrowhead), and basal cells spread flat on the insert membrane.

**Figure 1:**
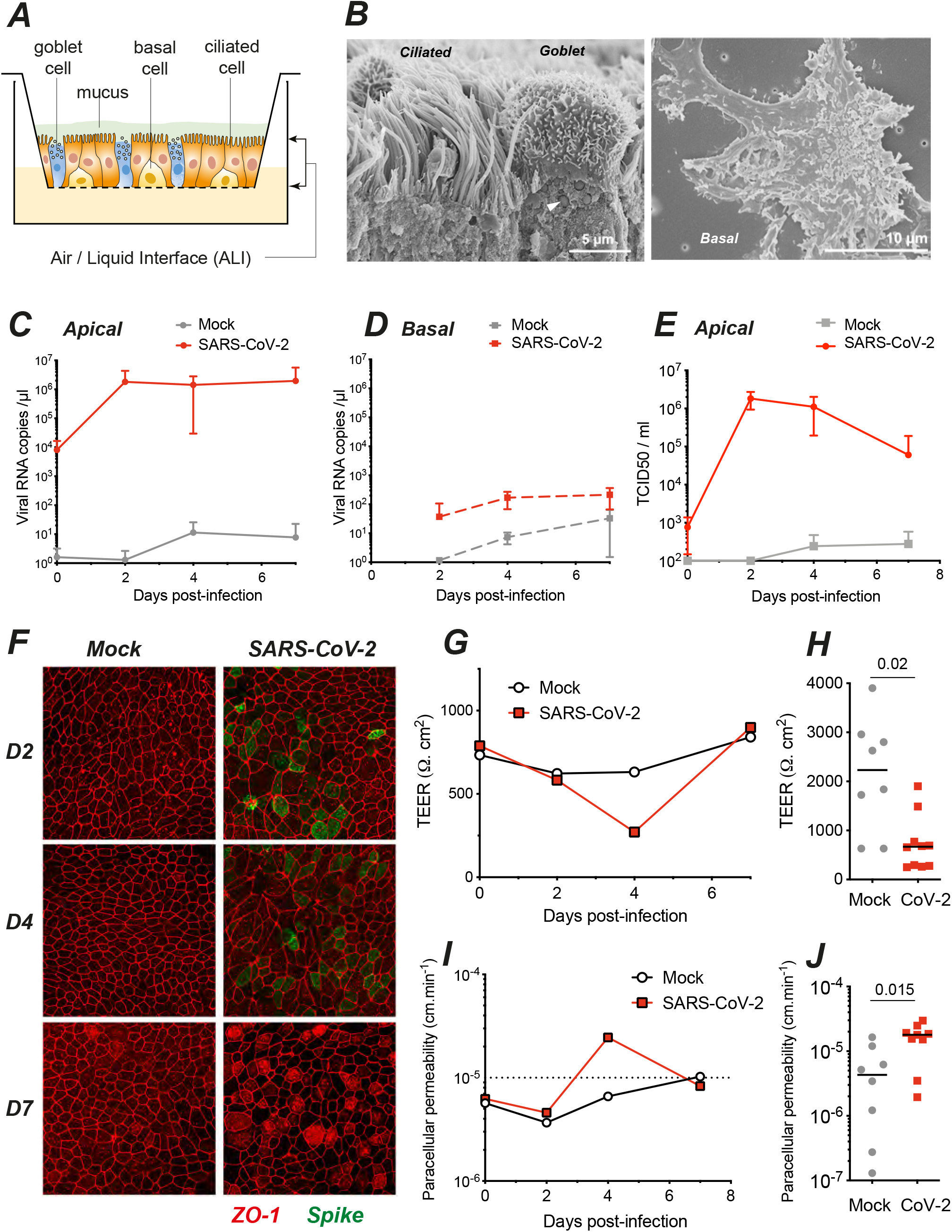
SARS-Cov-2 infection transiently impairs epithelial barrier function in a reconstituted human bronchial epithelium. **(A)** Schematic view of an Air/Liquid Interface (ALI) culture of a reconstituted human bronchial epithelium comprising three cell types: ciliated, goblet, and basal cells. **(B)** SEM imaging of the three cell types, with a ciliated and a goblet cell (left panel; arrowhead: mucus granule) and a basal cell (right panel). **(C-D)** SARS-CoV-2 viral load quantification in apical (C) and basal (D) culture supernatants by RT-qPCR, for n≥4 independent experiments (mean±SD). **(E)** Infectious SARS-CoV-2 particles production quantified by TCID_50_ on Vero cells (n≥5 replicates from 2 independent experiments; mean±SD). **(F)** Visualization of tight junctions (ZO-1) and infection (SARS-CoV-2 spike) by immunofluorescence at 2, 4, and 7 dpi. **(G-J)** Assessment of epithelium barrier function: Trans-Epithelial Electric Resistance (TEER) (G, H) and paracellular permeability for dextran-FITC (I, J). (G) and (I) show representative experiments, while (H) and (J) show pooled results at 4 dpi (n=4 and n=3 independent experiments, respectively). Median values were compared with Mann-Whitney tests.

The reconstituted epithelia were infected with a viral suspension of SARS-CoV-2 at 10^6^ pfu/mL deposited on the apical side for 4h, restored to ALI conditions, and monitored for infection for 7 days. We observed a rapid increase of extracellular viral RNA in apical culture supernatants, with concentrations reaching up to 10^6^ viral RNA copies/μL at 2 days post-infection (dpi), followed by stable or slowly decreasing viral RNA levels until 7 dpi (Fig. 1C). In contrast, minimal concentrations of viral RNA were detected in the basolateral compartment (Fig. 1D), indicating that SARS-CoV-2 particles were predominantly released from the apical side of the epithelium. Infectious viral particle production in apical supernatants initially tracked with viral RNA production, with a rapid increase at 2 dpi to reach a mean of 1.8 x 10^6^ TCID_50_/mL, followed by a plateau at 4 dpi (Fig. 1E). A decrease to 6.1 x 10^4^ TCID_50_/mL was then observed at 7 dpi, suggesting a partial containment of infectious virus production. Immunofluorescence analysis of the reconstituted epithelia revealed a patchy distribution of infected cells expressing the SARS-CoV-2 spike protein at 2 and 4 dpi, and minimal persistence of productively infected cells at 7 dpi, consistent with partial viral containment after one week of infection (Fig. 1F).

### SARS-CoV-2 infection transiently impairs epithelial barrier function

SARS-CoV-2 productive infection locally altered the distribution of zonula occludens protein-1 (ZO-1), which associates to tight junctions. The characteristic ZO-1 staining pattern at cell boundaries remained intact in mock-infected epithelia (Fig. 1F, left), but appeared disrupted in areas with viral antigen expression (Fig. 1F, right), suggesting a possible impairment of epithelial barrier integrity. Image analysis confirmed that the average number of neighbors per cell at 4 dpi decreased from 5.93 to 5.72 (P<0.001), a value characteristic of actively remodeling epithelia (Aigouy et al., 2010; Farhadifar et al., 2007) (Fig. S1A-H, J). In addition, infection induced changes in the average area of ZO-1-delimited cells, which significantly increased at 2 and 4 dpi, but decreased at 7 dpi as compared to uninfected samples (Fig.S1A-I), supporting a dynamic remodeling of the epithelium. The epithelial packing geometry returned to a more regular pattern at 7 dpi, with a higher proportion of hexagonal cells than at 4 dpi, pointing to the restoration of the tight junction network.

To test barrier function, we measured the trans-epithelial electrical resistance (TEER) between electrodes placed in the apical and basal compartments of reconstituted bronchial epithelia (Fig. 1G). As expected, TEER proved relatively stable in mock-treated cultures, with values remaining above 600 Ω.cm2. In SARS-CoV-2 infected cultures, there was a transient but significant 3.3x TEER decrease at 4 dpi (Fig. 1H; P=0.02). Apical-to-basolateral transport of FITC-coupled dextran showed an inverse relation to TEER, with a transient 2.5x increase in paracellular permeability at 4 dpi (Fig. 1I-J; P=0.015). Both TEER and permeability values returned to baseline levels at 7 dpi, demonstrating epithelial regeneration after SARS-CoV-2 infection.

Measurement of cell death by the release of lactate dehydrogenase (LDH) in apical supernatants showed an increase at 4 dpi in infected epithelia (Fig. S2A; P=0.01), confirming that SARS-CoV-2 exerted a transient cytopathic effect on epithelial cells. Dying cells extruded from the apical side were observed at the surface of infected epithelia, though their numbers remained limited (Fig. S2B). Extruded cells had a rounded shape and sometimes carried multiple viral particles at their surface (Fig. S2C), suggesting virally-induced cell death. Taken together, these findings showed that SARS-CoV-2 caused a transient loss of epithelial barrier function, due to cytopathic effect and a perturbation of the tight junction network. However, SARS-CoV-2 infection did not cause a general disruption of the epithelial layer and was compatible with rapid epithelial regeneration.

### SARS-CoV-2 infection damages the cilia layer

The nature of infected cells was characterized by immunofluorescence confocal imaging. At 2 dpi, the majority of cells expressing the SARS-CoV-2 spike antigen at their surface (spike+ cells) co-expressed the cilia marker β-tubulin IV, confirming that ciliated cells represented the main viral targets in this model (Fig. 2A). At 4 dpi, the majority of spike+ cells still expressed β-tubulin IV, though we noted the presence of infected cells with weak or absent β-tubulin IV staining (Fig. 2B). Basal cells expressing the cytokeratin-5 marker did not appear infected, with the exception of rare cells that had lost their basal localization (Fig. S3A). Of note, some spike-negative basal cells changed morphology and appeared raised through the pseudo-stratified epithelium in infected samples, suggesting an epithelial response to virally-induced damage (Fig. 2B). We did not detect infected cells expressing the goblet cell marker MUC5AC (Fig. S3B). However, further analyses by SEM documented viral budding from cells with multiple secretory pores that may represent rare infected goblet cells (Fig. S3C). Interestingly, viral production was also documented in cells with a transitional phenotype characterized by the presence of both motile cilia and abundant secretory vesicles (Fig. S3D). Taken together, these results showed a preferential tropism of SARS-CoV-2 for ciliated epithelial cells, with occasional infection of transitional and secretory cells.

**Figure 2:**
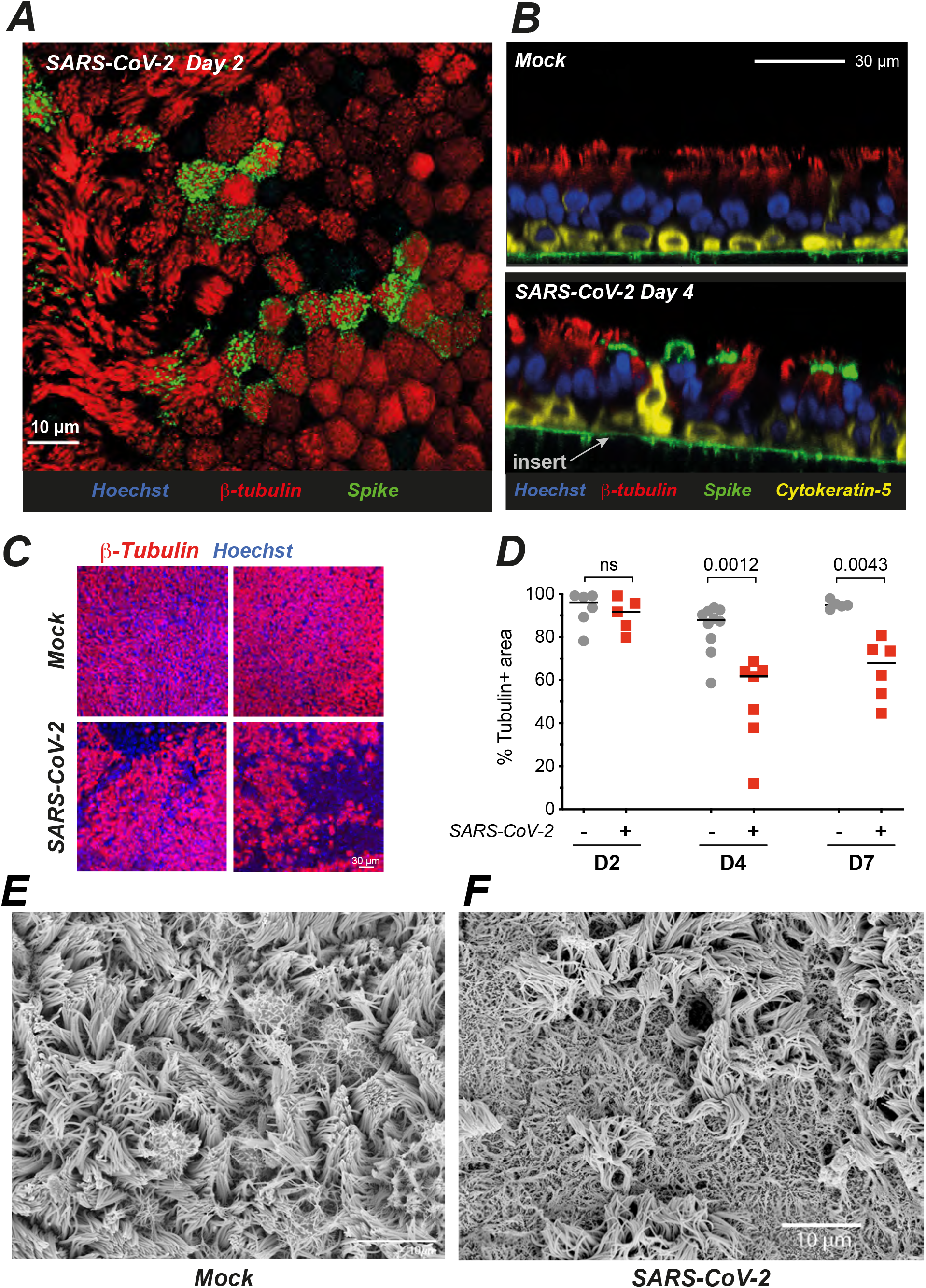
SARS-CoV-2 preferentially targets ciliated cells and damages the ciliary layer. **(A-B)** Confocal imaging of a SARS-CoV-2 infected epithelium at 2 dpi (A, top view) and of control (Mock) and infected epithelia at 4 dpi (B, orthogonal views). Ciliated cells are labeled for β-tubulin IV, basal cells for cytokeratin-5, nuclei for DNA (Hoechst), and infected cells for the SARS-CoV-2 spike. **(C-D)** Representative images of the β-tubulin layer at 4 dpi (C) and corresponding image analysis (D) (n=3 independent experiments; Mann-Whitney test). **(E-F)** SEM imaging of mock infected (E) and SARS-CoV-2 infected (F) reconstituted epithelia at 4 dpi.

As we had observed infected cells with weak β-tubulin IV staining, we asked whether SARS-CoV-2 infection could perturb the layer of motile cilia present at the apical side of ciliated cells. To this goal, we quantified the area occupied by β-tubulin IV staining (tubulin+ area) on projections of confocal images obtained sequentially in the course of infection (Fig. 2C-D). The tubulin+ area remained unchanged at 2 dpi, but decreased at 4 dpi (median values: 87.9% in mock, 61.7% in infected; P=0.0012), and showed only limited recovery at 7 dpi (94.8 % in mock vs 67.9% in infected, P=0.0043). SEM imaging confirmed a marked cilia loss at 4 dpi and showed that deciliated areas were not devoid of cells, but rather occupied by cells covered by flattened microvilli (Fig. 2E).

The formation of deciliated areas could occur via the loss of motile cilia at the surface of infected cells, or via the replacement of dead ciliated cells by cells involved in epithelial regeneration. To distinguish between these non-exclusive possibilities, we analyzed the distribution of cilia at the early stage of infection, prior to the occurrence of measurable cell death. A set of ≥70 cells was analyzed on confocal images obtained at 2 dpi in each of three categories: ciliated cells from mock-infected epithelia (mock), productively infected ciliated cells (spike+) and bystander uninfected ciliated cells (spike-) from SARS-CoV-2 exposed epithelia. For each cell, the averaged intensity profile of β-tubulin IV staining was measured along the depth axis (Fig. 3A). There was a bimodal distribution of β-tubulin in mock cells, with a distal peak corresponding to cilia and a proximal peak located just below the plasma membrane, corresponding to the area where basal bodies anchor cilia into the cytoplasm. Examination of average profiles for each category suggested a specific decrease of the distal β-tubulin peak in spike+ cells. This was confirmed by an analysis of the distal to proximal peak intensity ratio, which showed a significant decrease in spike+ cells, as compared to spike- and mock cells (Fig. 3B). Thus, the density of cilia decreased at 2 dpi, supporting an early loss of cilia in infected cells. SEM imaging confirmed the presence of productively infected cells with only few remaining cilia on their apical surface (Fig. 3C), as opposed to the packed ciliary layer characteristic of intact ciliated cells (Fig. 1B). Some infected cells showed a lack of cilia and a massive accumulation of virions at the cell surface and on membrane ruffles (Fig. 3D), indicative of highly productive SARS-CoV-2 infection. Of note, viral particles were rarely observed along the length of ciliary sheaths. This observation was consistent with the distribution of spike staining, which formed a narrow band above the plasma membrane, but did not overlay the distal β-tubulin peak (Fig. 3A, spike+; Fig. S4A). In addition, spike and β-tubulin labeling showed minimal colocalization, as measured by Mander correlation coefficients (Fig. S4B-C). Thus, viral budding or accumulation did not take place in the cilium structure, suggesting that cilia destruction occurred through an indirect mechanism.

**Figure 3:**
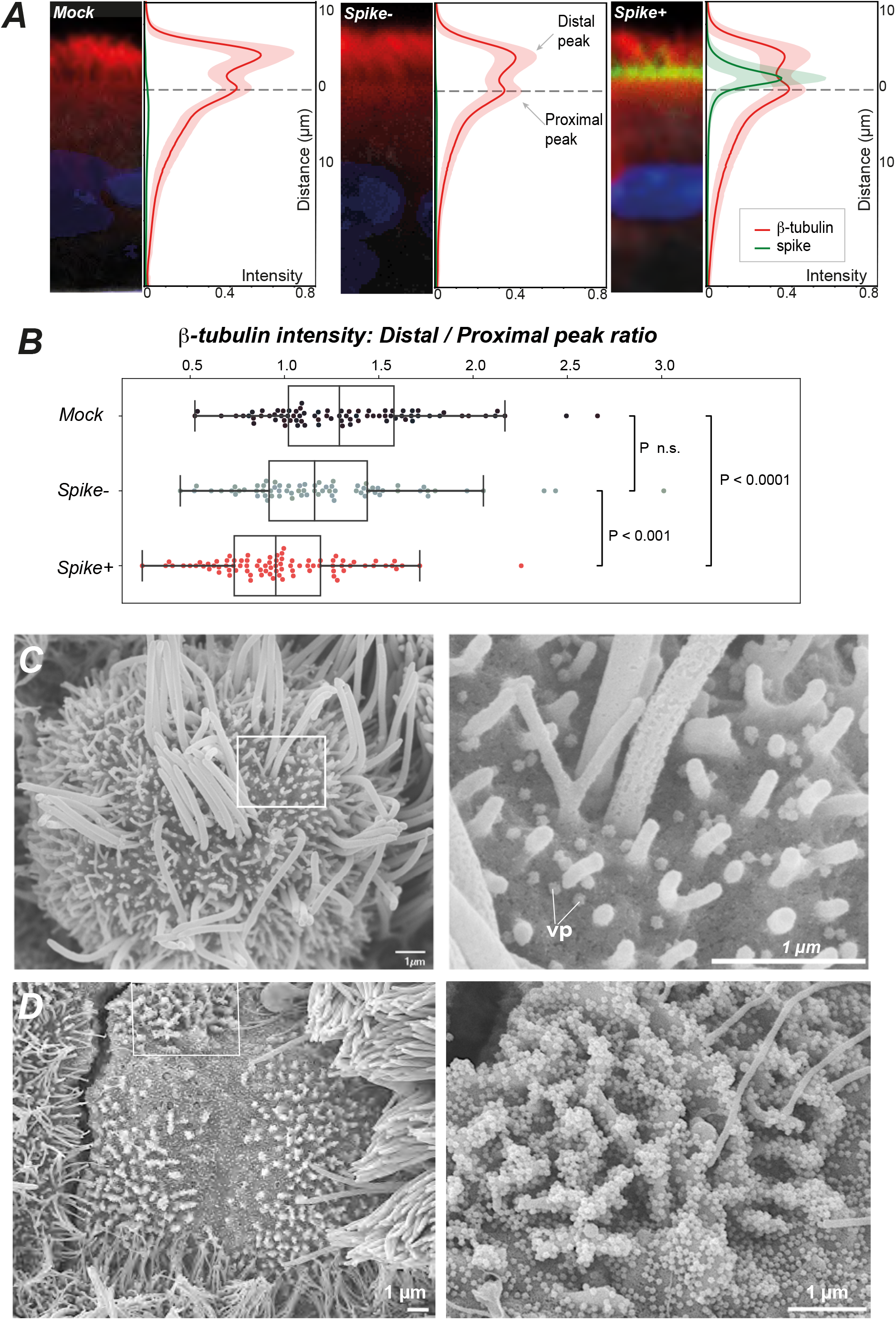
SARS-CoV-2 infection causes an early loss of motile cilia. **(A)** Distribution of β-tubulin IV and SARS-CoV-2 spike immunofluorescence (IF) labeling at 2 dpi. Representative IF images projected on the Z (depth) axis are shown next to averaged profiles of β-tubulin intensities. Profiles were measured along the Z axis for n≥70 cells (mean±SD) from uninfected (mock) and infected (spike+, spike-) epithelia. **(B)** Ratio of β-tubulin distal to proximal peak intensities for 3 cell categories. **(C)** SEM image of an infected cell at 2 dpi with few remaining cilia (left) and scattered viral particles (vp) at the plasma membrane (right). **(D)** SEM image of a massively infected cell at 2 dpi (left) with an accumulation of vp at the surface of membrane ruffles (right). (C, D) Framed regions in left panels are enlarged in right panels.

### SARS-CoV-2 infection induces ultrastructurally abnormal cilia

SEM imaging performed at higher magnification revealed ultrastructural abnormalities in infected ciliated cells. Cilia were often shortened and misshapen (Fig. 4A-B) and sometimes showed crescent-shaped juxta-membrane regions (Fig. 4C). Viral particles present at the membrane were not symmetrical but rather pleiomorphic (Fig. 4D), consistent with studies suggesting that spike trimers adopt various orientations at the virion surface (Turonova et al., 2020). Transmission electron microcopy (TEM) was then used to examine the internal organization of cilia. Mock-infected epithelia revealed a typical “9+2” cilium structure, with peripheral microtubule doublets surrounding a central microtubule pair (Fig. S4D). These microtubules constituted elongated axonemes that emerged from basal bodies aligned perpendicular to the plasma membrane (Fig. 4E). The basal bodies were themselves anchored into the cytoplasm through striated rootlets (Fig. 4F). A striking disorganization of cilia structure was observed in SARS-CoV-2 infected cells, with fewer axonemes, and misoriented basal bodies that lined large vesicles, which themselves often contained viral-like particles (Fig. 4G and S4E). Isolated rootlets were also detected, suggesting a dissociation of ciliary components (Fig. S4G). The vesicles contained packed viral-like particles, but also larger particles that may have derived from engulfed basal bodies (Fig. 4H and S4E). Therefore, SARS-CoV-2 infection had a major impact on ciliary structure, by inducing axoneme loss and accumulation of mislocalized basal bodies.

**Figure 4:**
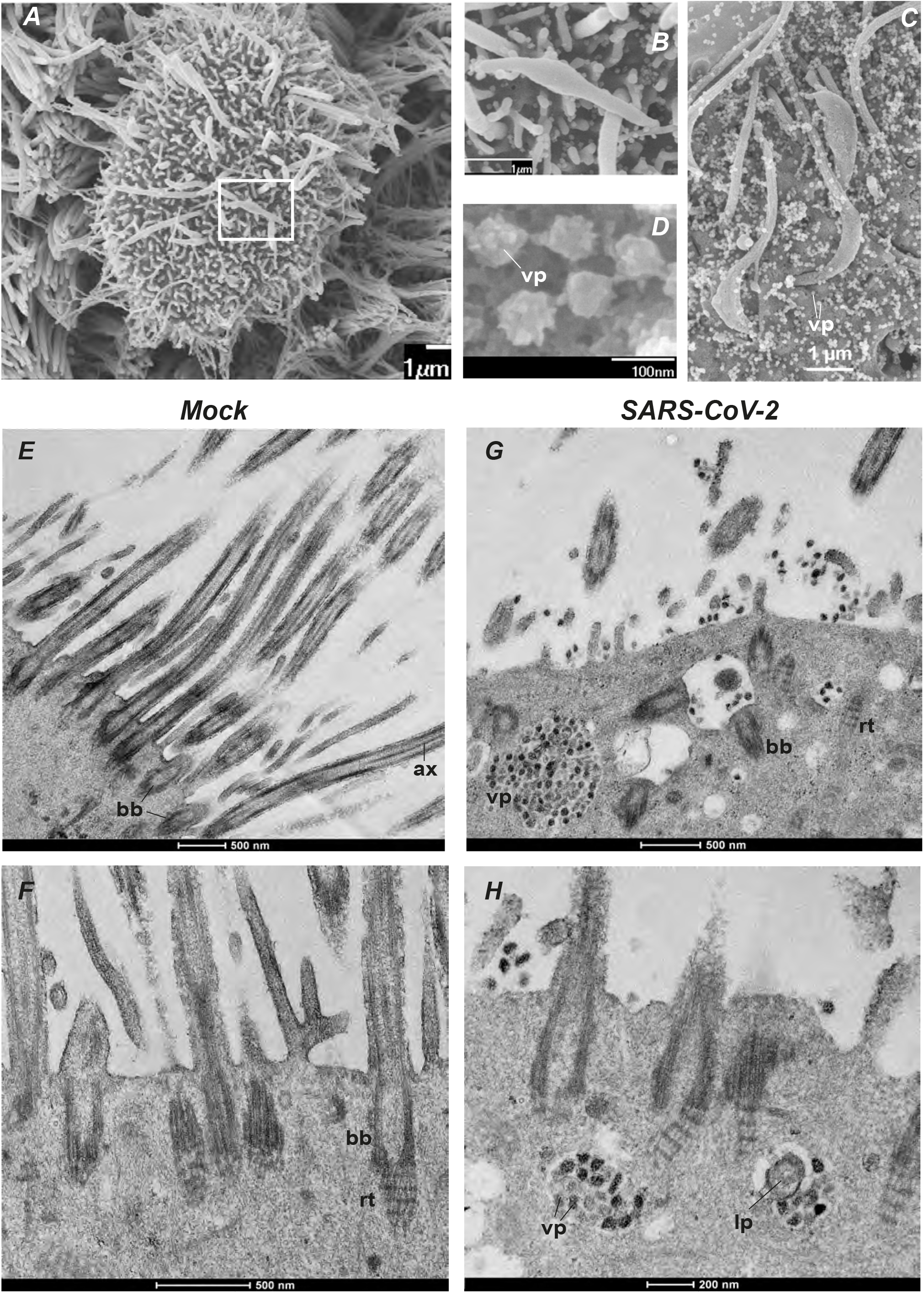
SARS-CoV-2 infection induces ultrastructurally abnormal cilia. **(A-C)** SEM images of infected cells at 2 dpi showing cilia abnormalities, including cilia loss and shortened cilia (A; enlarged in B), and cilia with crescent-shaped proximal axonemes (C). **(D)** SEM image of SARS-CoV-2 viral particles. **(E-H)** TEM images of mock (E-F) and infected (G-H) ciliated cells at 4 dpi. bb: basal body; ax: axoneme; rt: rootlet; vp: viral particles; lp: large particle.

### SARS-CoV-2 infection impairs mucociliary clearance

We next examined the consequences of SARS-CoV-2 infection on ciliary function. To this goal, we deposited low density 30 μm-sized polystyrene microbeads onto the apical surface of mock-treated or infected epithelia and tracked their movement in real time. These experiments were performed at 7 dpi, to allow sufficient reconstitution of the mucus layer after the infection step. We observed that beads deposited on mock-treated epithelia moved generally in the same direction, consistent with coordinated beating of the underlying cilia (Fig. 5A, 5C, and Supplemental Movie 1). In contrast, beads deposited on infected epithelia were mostly immobile or showed randomly-oriented limited movements, indicating an impairment of the mucociliary clearance function (Fig. 5B, 5D, and Supplemental Movie 2). Quantitation of velocities confirmed a highly significant decrease in bead clearance upon SARS-CoV-2 infection (mean: 8.9 μm/s in mock vs 1.5 μm/s in infected epithelia; P<0.0001) (Fig. 5E). The straightness of tracks also decreased in infected epithelia (Fig. 5F; P<0.0001), suggesting a perturbation in the coordination of cilia movements. Thus, cilia alterations induced by SARS-CoV-2 infection were associated to a marked impairment in mucociliary transport.

**Figure 5:**
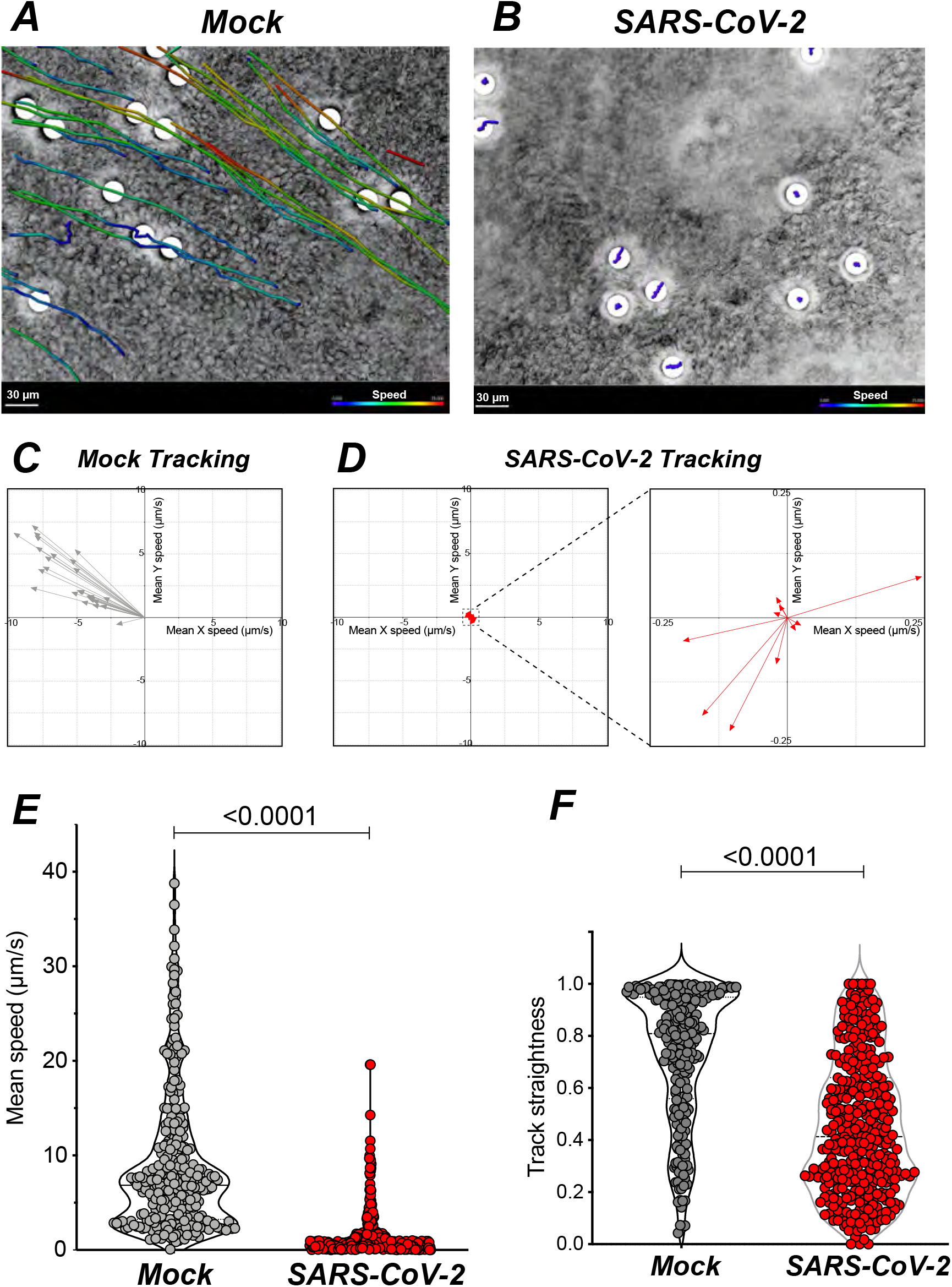
SARS-CoV-2 infection impairs mucociliary clearance. **(A-B)** Representative examples of bead tracking on mock (A) and infected (B) epithelia at 7 dpi. Bead speed is color-coded in each track. **(C-D)** Vector representation of tracks from (A) and (B) showing bead speed and direction. A low bead speed is observed in infected epithelia (D, left), with an enlarged view showing non-uniform directions (D, right). **(E-F)** Bead mean speed (E) and track straightness (F) measured in mock and infected epithelia at 7 dpi (n=3 independent experiments; Mann Whitney tests).

### Epithelial defense mechanisms are triggered by SARS-CoV-2 infection

Mucociliary transport was altered at 7 dpi, at a time when barrier integrity was already restored (Fig. 1). To better apprehend the process of epithelial regeneration, we analyzed the localization and morphology of basal cells at this time point. Confocal images showed that basal cells expressing cytokeratin-5 were typically flattened on the basement membrane of mock-treated epithelia, while they appeared raised through the thickness of the pseudo-stratified epithelium in infected samples (Fig. 6A). To quantify this phenomenon, we first generated elevation maps of the inserts that support the cultures, and used these to correct for local insert deformations (Fig. S5A). This approach enabled to precisely quantify the mean height of basal cells in the epithelia, which proved significantly higher in infected than non-infected samples (Fig. 6B-C). SEM imaging confirmed that basal cells adopted a more rounded morphology in infected epithelia (Fig. S5B). Thus, basal cells were mobilized at 7 dpi, which may contribute to the restoration of barrier integrity. However, they had not differentiated into ciliary cells at this stage, as attested by the persistence of deciliated areas (Fig. 2D) and the impairment in mucociliary clearance (Fig. 5E).

**Figure 6:**
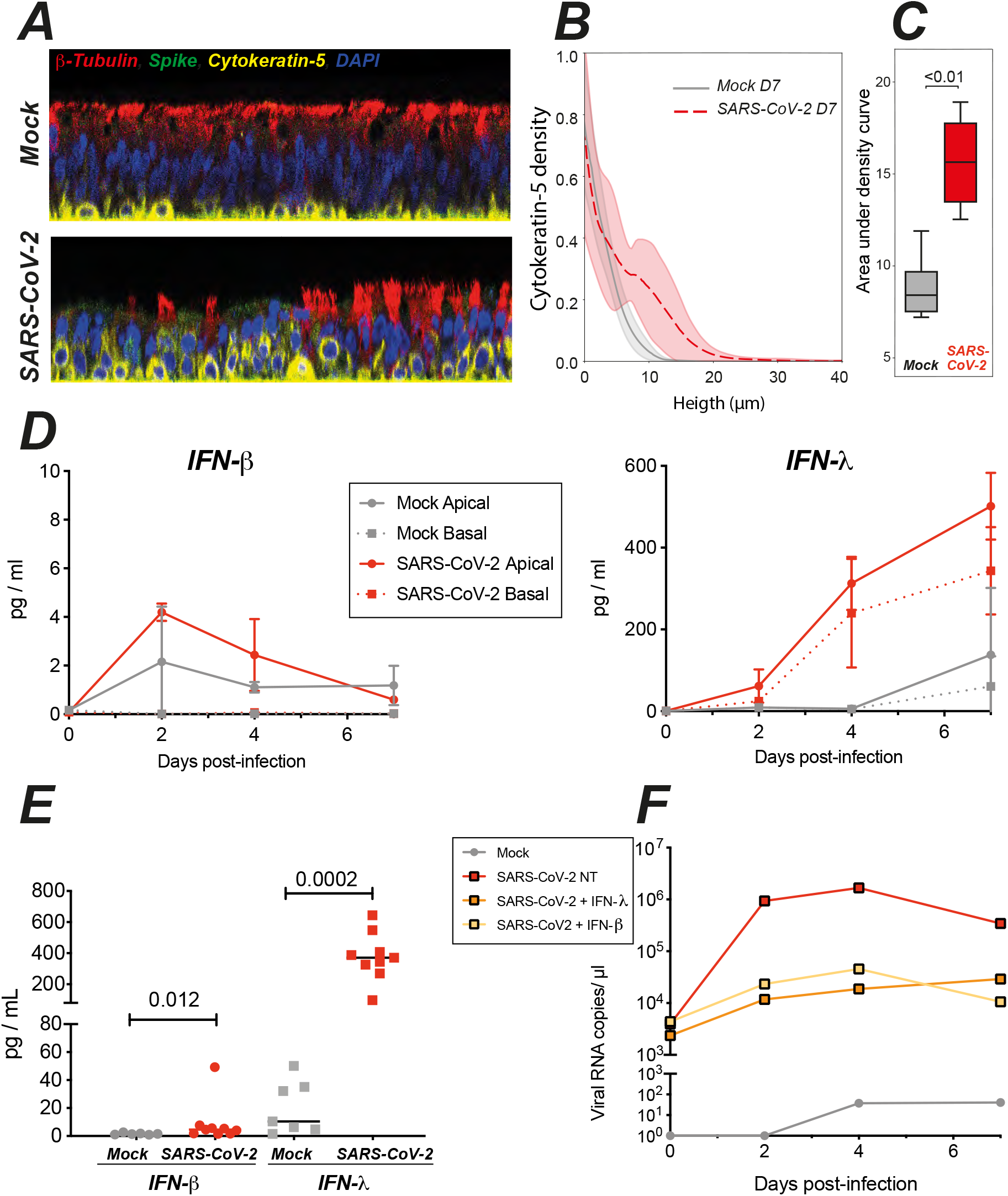
SARS-CoV-2 infection triggers epithelial defense mechanisms: basal cell mobilization and interferon induction. **(A)** Representative confocal images from mock and infected epithelia obtained at 7 dpi and used for quantification of basal cell (cytokeratin-5+) position along the Z axis. **(B-C)** Quantification of the cytokeratin-5 density profile along the Z axis. (B) The mean density +/- SD of n=9 images in each group is shown. (C) Comparison area under the curves shown in (B) with a Welch t-test. **(D)** Kinetics of IFN-β (left) and IFN-λ (right) production in apical and basal supernatants of mock and infected epithelia (n=2 independent experiments). **(E)** Quantification of interferon production in apical supernatants at 4 dpi (n=4 independent experiments; Mann Whitney test). **(F)** Effect of interferon pretreatment on SARS-CoV-2 replication in reconstituted epithelia. Viral RNA copies were quantified in apical supernatants by qRT-PCR (one representative experiment).

Epithelial interferon production provides another key defense mechanism, that can occur in the absence of immune cell infiltration, and thus represents one of the earliest antiviral response to respiratory viruses. We monitored the kinetics of type I (IFN-a, IFN-β) and type III (IFN-λ) interferon production in the supernatants of reconstituted epithelia. IFN-α⍰ remained undetectable (data not shown), using a high-sensitivity SIMOA assay with a limit of detection (LOD) of 2 fg/mL (Hadjadj et al., 2020). IFN-β was minimally induced upon SARS-CoV-2 infection, with peak values of 4.2±0.4 pg/mL at 2 dpi in apical supernatants, and levels below the LOD (< 1.7 pg/mL) in basal supernatants (Fig. 6D, left). In contrast, IFN-λ induction showed a different kinetics, with limited induction in apical supernatants at 2 dpi, but persistent increase at 4 and 7 dpi, to reach relatively high concentrations in both the apical (501.3±81.8 pg/mL) and basal supernatants (343±106 pg/mL) (Fig. 6D, right). Both IFN-β and IFN-λ production showed a degree of inter-sample variability, but were significantly induced as compared to mock-treated samples at 4 dpi (Fig. 6E). It remained striking, however, that viral replication already peaked at 2 dpi (Fig. 1E), while interferon production was minimal at this stage. Pretreatment of the reconstituted epithelia with exogenous IFN-β or IFN-λ prior to SARS-CoV-2 infection decreased viral RNA levels by 2 logs (Fig. 6F), pointing to the importance of the timing of IFN induction to achieve viral containment. Taken together, these findings documented the induction of epithelial defense mechanisms following SARS-CoV-2 infection, including basal cell mobilization and type III interferon induction. However, the kinetics of these responses appeared too slow in this model to prevent viral replication and functional impairment.

### SARS-CoV-2 infection damages the ciliary layer in the respiratory tract of Syrian hamsters

We next asked whether the ciliated airway epithelium is impacted by SARS-CoV-2 infection *in vivo*. We chose the golden Syrian hamster (*Mesocricetus auratus*) model, as this species is naturally susceptible to SARS-CoV-2 and shows lung lesions upon infection (Chan et al., 2020; Sia et al., 2020). Syrian hamsters were infected with 6 x 10^4^ pfu of SARS-CoV-2 via the intranasal route. The infected hamsters showed rapid body weight loss as compared to mock-infected controls (n=4 in each group; Fig. 7A). The animals were euthanized at 4 dpi, at a time when the infected group showed a high viral load in the trachea (Fig. 7B). SEM imaging showed that cilia occupied almost half of the epithelial surface in the trachea of control animals (Fig. 7C, left). A marked cilia loss occurred in the trachea of infected animals (Fig. 7C, right), as confirmed by image analysis (median of ciliated area: 2.5% in SARS-CoV-2+ vs 45.4% in Mock, P=0.0079) (Fig. 7D). Immunofluorescence labeling of trachea sections confirmed a partial to complete loss of the ciliated layer at 4 dpi, with a mobilization of basal cells towards the luminal side of the epithelium (Fig. 7E). The SARS-CoV-2 spike antigen could be detected in a few remaining ciliated cells (Fig. S6). Therefore, SARS-CoV-2 infected and damaged the epithelial ciliary layer in a physiologically relevant animal model.

**Figure 7:**
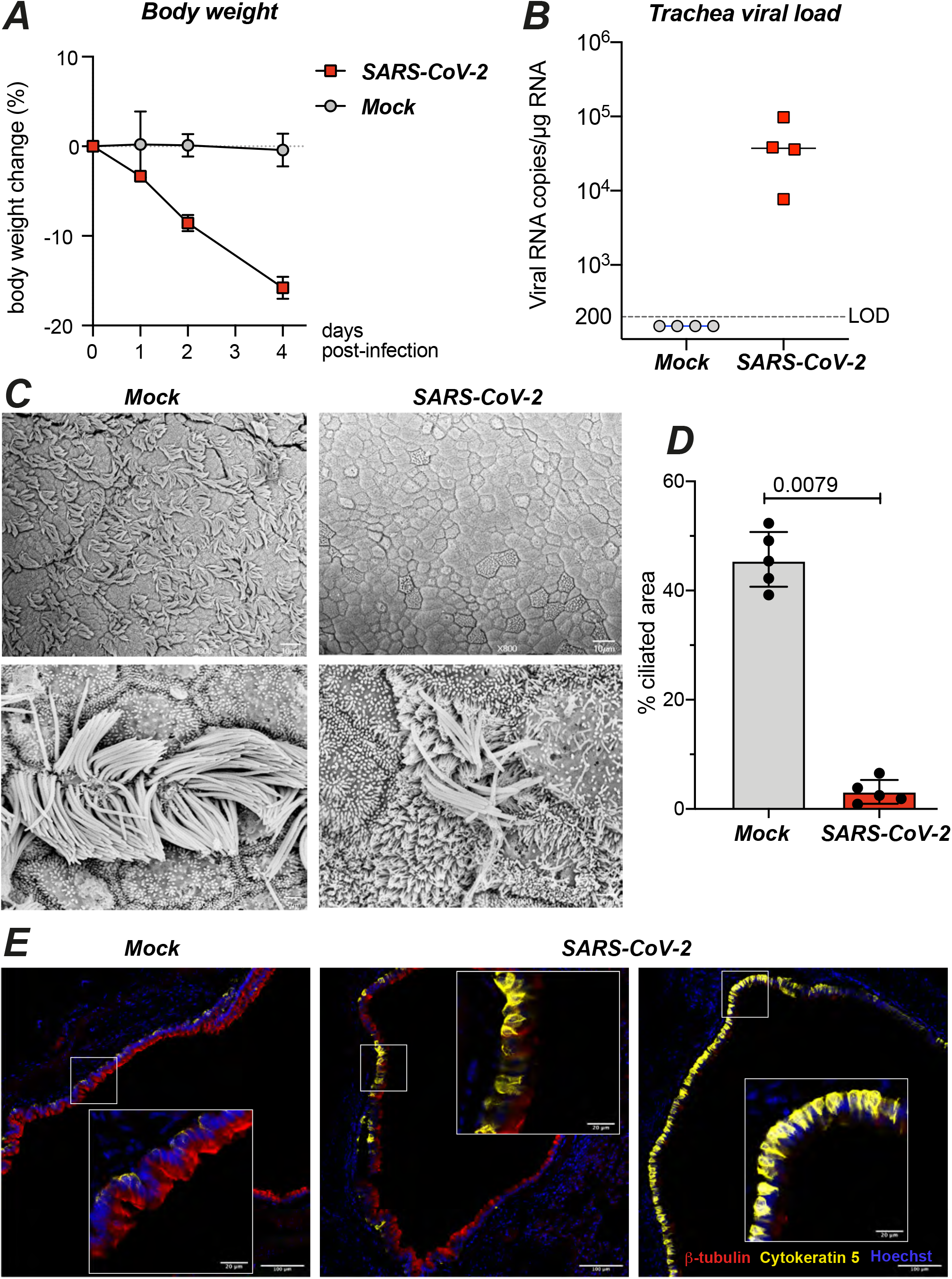
SARS-CoV-2 damages the ciliary layer in the respiratory tract of Syrian hamsters. **(A)** Effect of intranasal SARS-CoV-2 infection on hamster body weight (n=4 per group). **(B)** SARS-CoV-2 viral load measured at 4 dpi in hamster trachea by RT-qPCR (n=4 per group). **(C)** SEM images of tracheal epithelia from mock (left panels) and SARS-CoV-2 infected (right panels) hamsters at 4 dpi. Enlarged views of ciliated cell are shown in bottom panels. **(D)** Quantification of ciliated area in hamster tracheas imaged by SEM (n=5 images per group). Differences between medians ±IQR are measured by a Mann Whitney test. **(E)** IF imaging of hamster tracheas at 4 dpi. Ciliated cells are labeled with β-tubulin IV, basal cells by cytokeratin-5, and nuclei by Hoechst. Representative images are shown for a mock infected animal (left), and SARS-CoV-2 infected animals who received a low dose (middle) or a high dose (right) of virus.

## DISCUSSION

We report here that SARS-CoV-2 preferentially replicates in ciliated cells, damages the ciliary layer, and impairs mucociliary clearance in a reconstituted human bronchial epithelium model. We also demonstrate, in the SARS-CoV-2 susceptible hamster model, a loss of motile cilia in the trachea of infected animals. Thus, SARS-CoV-2 specifically damaged airway motile cilia, both *in vivo* and *in vitro*. Our findings suggest that cilia loss may also play a role in COVID-19 pathogenesis, as a localized clearance impairment at the site of SARS-CoV-2 replication could facilitate viral spread within the airways. Decreased cilia movement could slow the transport of released virions towards the pharynx and facilitate viral access to deeper regions of the bronchial tree. This process could self-perpetuate, with cycles of localized cilia destruction facilitating SARS-CoV-2 progression towards increasingly more distal regions, until the virus reaches the alveoli and triggers pneumocyte damage.

We did not observe major discontinuities in the tracheal epithelium of infected hamsters. Continuity of the epithelial layer was not disrupted in the *in vitro* model either, as confirmed by the minimal release of viral particles in the basal compartment. A degree of cytotoxicity was measured by LDH release, but the observation of extruded dead cells remained occasional. Epithelial barrier function was impaired at day 4, as documented by decreased transepithelial resistance, increased permeability, and altered distribution of the tight junction protein ZO-1. However, this functional impairment remained transient, with signs of epithelial regeneration such as basal cell mobilization observed at day 7. Our results indicate that SARS-CoV-2 replication has a drastic effect on the ciliary layer while exerting a moderate effect on epithelial barrier integrity. This is in contrast to other respiratory viruses such as influenza A virus (IAV), which induces widespread epithelial cell death and persistent loss of epithelial barrier function (Essaidi-Laziosi et al., 2018). Respiratory viruses use distinct strategies that all converge in limiting mucociliary clearance, including motile cilia loss (this study)(Look et al., 2001), epithelial disruption (Essaidi-Laziosi et al., 2018; Villenave et al., 2010), and altered ciliary movements (Essaidi-Laziosi et al., 2018; Smith et al., 2014). Experimental impairment of ciliary movements through chemical treatment also increased IAV infection *in vitro* (Fu et al., 2018), providing further evidence that decreasing mucociliary clearance can facilitate viral infection.

The role of mucociliary clearance in limiting respiratory bacterial infections is also well documented (Gudis et al., 2012), and supported by the recurrence of severe pulmonary infections in patients with inborn errors in genes essential to motile cilia function (Shapiro et al., 2016). The percentage of COVID-19 patients presenting with a bacterial or fungal respiratory co-infection at hospital admission remains moderate (Garcia-Vidal et al., 2020; Rawson et al., 2020). However, the occurrence of secondary infections increases in critically-ill COVID-19 patients, in spite of common broad-spectrum antibiotic prophylaxis. One fourth to one third of severe COVID-19 patients experience secondary bacterial or fungal superinfections (Alanio et al., 2020; Manohar et al., 2020; White et al., 2020). Secondary infections are associated with worse outcomes (Garcia-Vidal et al., 2020; Manohar et al., 2020), emphasizing the possible effect of impaired mucociliary clearance on the risk of severe COVID.

Ultrastructural analysis by SEM revealed a loss of motile cilia in productively infected human bronchial cells. Misshapen and shortened cilia were also detected, reminiscent of abnormalities observed in certain cases of primary ciliopathies (Shah et al., 2008; Tilley et al., 2015). We documented by immunofluorescence a thinning of the ciliary layer as early as 2 dpi, associated to a layer of viral spike protein at the base of motile cilia. SARS-CoV-2 particles did not bud from cilia, but rather from deciliated areas, as indicated by the minimal colocalization between the spike and β-tubulin markers. Clusters of viral particles could be detected on microvilli, consistent with findings suggesting that SARS-CoV-2 could induce the formation of actin-based filaments in transformed cells (Bouhaddou et al., 2020). The absence of viral particles within cilia may result from the restricted protein access imposed by the transition zone at the base of motile cilia (Nachury and Mick, 2019). Access of cellular proteins into the ciliary axoneme is limited to those bound by the intraflagellar transport machinery, which likely prevents the import of viral proteins. Therefore, the destruction of motile cilia by SARS-CoV-2 seems mediated by an indirect mechanism, rather than by direct viral production within these structures. TEM imaging revealed the presence of misoriented basal bodies that had lost plasma membrane docking in productively infected cells. The ciliary axoneme is normally maintained through a dynamic equilibrium between microtubule polymerization/depolymerization, in turn regulated by the supply of ciliary components by intraflagellar transport (Lechtreck et al., 2017). Our observations in infected cells suggest a disequilibrium favoring depolymerization, leading to loss of ciliary axonemes. Rootlets, which normally anchor basal bodies into the cytoplasm, were sometimes found isolated, suggesting that basal bodies may themselves detach and/or depolymerize. It is relevant that RNAseq studies of SARS-CoV-2 infected epithelial cell cultures (Nunnari et al., 2020; Ravindra et al., 2020) and of mucosal samples from COVID-19 patients (He et al., 2020) reported a downregulation of transcripts involved in cilia function, such as dyneins, supporting the idea of a virally-induced dedifferentiation of ciliated cells. Future work will help decipher the step at which SARS-CoV-2 impairs the production, stability, or transport of ciliary components.

SARS-CoV-2 infection also induced a significant IFN response, that was primarily driven by type III IFN. IFN-λ was abundantly secreted at 4 dpi in the apical and basal compartments of infected reconstituted epithelia. In contrast, IFN-β secretion was minimal in the apical compartment and undetectable in the basal compartment. IFN-a secretion remained undetectable in both compartments, even though we used a high-sensitivity Simoa digital ELISA for detection. These observations fit with the notion that type III IFN is the dominant antiviral cytokine in epithelia, enabling a localized response mediated directly by epithelial cells, prior to the infiltration of immune cells (Park and Iwasaki, 2020). It was noteworthy, however, that IFN was secreted mostly from day 4 post-infection onward, while viral RNA production had already shown a 2 log increase by day 2. This delay in the induction of the antiviral response suggests a “too little too late” scenario, where antiviral mechanisms are overwhelmed and subverted by active viral replication. Our findings are compatible with studies showing a limited and/or delayed IFN induction by SARS-CoV-2 as compared to other respiratory viruses, in primary epithelial cells from the airways and the intestine (Blanco-Melo et al., 2020; Lamers et al., 2020; V’kovski et al., 2020; Vanderheiden et al., 2020).

These results do not rule out the potential for IFN treatment, as IFN may still exert an antiviral effect if administered early. We observed that pretreatment with IFN-β or IFN-λ markedly decreased SARS-CoV-2 replication in our reconstituted bronchial epithelium model. These findings are in agreement with studies in various culture systems documenting the inhibitory effect of both type I and type III IFN on SARS-CoV-2 replication, when treatment occurs prior to the stage of massive viral replication (Felgenhauer et al., 2020; Mantlo et al., 2020; Stanifer et al., 2020; V’kovski et al., 2020; Vanderheiden et al., 2020). However, studies in animal models suggest that, at an advanced stage of infection, IFN may contribute to the decline of respiratory function, possibly by worsening inflammation (Boudewijns et al., 2020) and impairing epithelial cell regeneration (Broggi et al., 2020; Major et al., 2020). Therefore, the window of opportunity for IFN treatment may have to be carefully defined in randomized controlled trials.

In conclusion, we demonstrate that SARS-CoV-2 induces a rapid loss of the ciliary layer in airway epithelial cells, resulting in impaired mucociliary clearance. Infected cells display ultrastructurally altered motile cilia, characterized by misshapen or absent axonemes. Intrinsic epithelial defense mechanisms likely occur too late to prevent cilia loss. These findings highlight a pathogenic mechanism that may promote viral spread in the respiratory tree and increase the risk of secondary infections in COVID-19 patients.

## ACKNOWLEDGEMENTS

We thank Charlotte Calvet for help with sample preparation for scanning electron microscopy, Sylvie Van der Werf for the SARS-CoV-2 isolate used in this study, and Nicolas Escriou for the gift of a SARS-CoV-2 spike antibody.

This work was supported by: Institut Pasteur TASK FORCE SARS COV2 (Tropicoro project), DIM ELICIT Region Ile-de-France, and ANRS (L.A.C.); the Vaccine Research Institute (ANR-10–LABX–77), ANRS, Labex IBEID (ANR-10-LABX-62-IBEID), “TIMTAMDEN” ANR-14-CE14-0029, “CHIKV-Viro-Immuno” ANR-14-CE14-0015-01, the Gilead HIV cure program, ANR/FRM Flash Covid PROTEO-SARS-CoV-2 and IDISCOVR (O.S.); Institut Pasteur TASK FORCE SARS COV2 and ANR Flash Covid CoVarImm (D.D.); Institut Pasteur TASK FORCE SARS COV2 (Neuro-Covid project) (H.B.). The Lledo’s lab is supported by the life insurance company “AG2R-La-Mondiale”. The UtechS Photonic BioImaging (Imagopole) and the UtechS Ultrastructural BioImaging (UBI) are supported by the French National Research Agency (France BioImaging; ANR-10-INSB-04; Investments for the Future). R.R. is the recipient of a Sidaction fellowship, N.S. of a Pasteur-Roux-Cantarini fellowship, and St.G. of a MESR/Ecole Doctorale B3MI, Paris 7 University fellowship. S.L. is supported by FRM (fellowship ECO201906009119) and by “Ecole Doctorale FIRE – Programme Bettencourt”.

## AUTHOR CONTRIBUTIONS

R.R., M.H., Sa.G., P.M.L., M.L., H.B., D.D., V.M., O.S., and L.A.C. designed the project

R.R., M.H., G.D.d.M., F.L.S., T.B., N.S., S.L., F.L., J.F., St.G., O.G., C.T., F.G.B., and V.M. performed experiments

R.R., T.B., J.F., S.R., C.T., A.M., G.D., R.E., and L.A.C. contributed to image analysis

R.R., M.H., V.M., O.S., and L.A.C. wrote the manuscript

All authors reviewed and approved the manuscript.

## DECLARATION OF INTERESTS

The authors declare that they have no competing interests.

## Notes

### Competing Interest Statement

The authors have declared no competing interest.

