## Supplemental text and figures for "SARS-CoV-2 infection damages airway motile cilia and impairs mucociliary clearance"

**CONTENT:**

**Methods**

**Supplemental references**

**Legends to the supplemental figures**

**Supplemental figures:**

Figure S1: Quantitative image analysis of the ZO-1-associated tight junction pattern

Figure S2: SARS-CoV-2 induces a cytopathic effect in reconstituted human bronchial epithelia

Figure S3: SARS-CoV-2 occasionally infects basal and secretory cells

Figure S4: Image analysis of cilia

Figure S5: Image analysis of basal cells

Figure S6: SARS-CoV-2 infection targets ciliated cells in the trachea of infected Syrian hamsters

### METHODS

### 1- EXPERIMENTAL MODELS

#### SARS-CoV-2 infection of reconstituted human bronchial epithelia

MucilAir$^{TM}$, corresponding to reconstituted human bronchial epithelium cultures differentiated in vitro for at least 4 weeks, were purchased from Epithelix (Saint_Julien_en_Genevois, France). Cultures were maintained in air/liquid interface (ALI) conditions in transwells with 700 µL of MucilAir$^{TM}$ medium (Epithelix) in the basal compartment, and kept at 37°C under a 5% CO2 atmosphere.

For SARS-CoV-2 infection, the apical side of ALI cultures was washed 20 min at 37°C in Mucilair$^{TM}$ medium to remove mucus. Cells were then incubated with $10^6$ plaque-forming units (pfu) of the isolate BetaCoV/France/IDF00372/2020 (EVAg collection, Ref-SKU: 014V-03890; kindly provided by S. Van der Werf). The viral input was diluted in DMEM medium to a final volume 150 µL, and left on the apical side for 4 h at 37°C. Control wells were mock-treated with DMEM medium (Gibco) for the same duration. Viral inputs were removed by washing twice with 200 µL of PBS (5 min at 37°C) and once with 200 µL Mucilair$^{TM}$ medium (20 min at 37°C). The basal medium was replaced every 2-3 days. Apical supernatants were harvested every 2-3 days by adding 200 µL of Mucilair$^{TM}$ medium on the apical side, with an incubation of 20 min at 37°C prior to collection.

For viral inhibition by interferons, cultures were treated with IFN-$\beta$ at 1000 U/mL or with a mixture of IFN-$\lambda$1 and IFN-$\lambda$3 at 1 µg/mL each. Interferons were added to the basal compartment 1 day prior to infection, and then added every 2-3 days to both the apical and basal compartments throughout the infection.

#### SARS-CoV-2 infection of Syrian hamsters

Male Syrian hamsters (*Mesocricetus auratus*) of 5-6 weeks of age (average weight 60-80 grams) were purchased from Janvier Laboratories and handled under specific pathogen-free conditions, according to the French legislation and to the regulations of Pasteur Institute Animal Care Committees, in compliance with the European Communities Council Directives (2010/63/UE, French Law 2013–118, February 6, 2013). The Animal Experimentation Ethics Committee (CETEA 89) of the Pasteur Institute approved this study (2020–0023). The animals were housed and manipulated in isolators in a Biosafety level-3 facility, with ad libitum access to water and food. Before any manipulation, animals underwent an acclimation period of one week.

Animal infections were performed as described with few modifications (Chan et al., 2020). Briefly, the animals were anesthetized with an intraperitoneal injection of ketamine (200 mg/kg) and xylazine (10 mg/kg). 100 µL of physiological solution containing $6x10^4$ pfu of SARS-CoV-2 (BetaCoV/France/IDF00372/2020) were administered intranasally. Mock-infected animals received the physiological solution only. The animals were followed-up on a daily basis and euthanized at day 4 post-infection, when the tracheas were collected, immediately frozen at -80°C or formalin-fixed after transcardial perfusion with a physiological solution containing heparin ($5x10^3$ U/ml, Sanofi), followed by 4% paraformaldehyde (PFA).

**2- METHOD DETAILS**

**Viral RNA quantification**

Reconstituted bronchial epithelia: viral RNAs were extracted from 20 µL of apical and basal culture supernatants using the Quick-RNA Viral 96 kit (Zymo) following manufacturer instructions. SARS-CoV-2 RNA was quantified in a final volume of 5 µL per reaction in 384-well plates using the Luna Universal Probe One-Step RT-qPCR Kit (New England Biolabs) with SARS-CoV-2 N-specific primers (Forward 5'-TAA TCA GAC AAG GAA CTG ATT A-3'; Reverse 5'-CGA AGG TGT GAC TTC CAT G-3') on a QuantStudio 6 Flex thermocycler (Applied Biosystems). Standard curve was performed in parallel using purified SARS-CoV-2 viral RNA.

Hamster tissues: total RNA was extracted from frozen tracheas using the Direct-zol RNA MicroPrep Kit (R2062, Zymo Research) and reverse transcribed to first strand cDNA using the SuperScript™ IV VILO™ Master Mix (Invitrogen). qPCR was performed in a final volume of 20 µL per reaction in 96-well PCR plates using a thermocycler (7500t real time PCR system, Applied Biosystems). Briefly, 5 µL of cDNA (25 ng) was added to 15 µL of a master mix containing 10 µL of Power SYBR green mix (4367659, Applied Biosystems) and 5 µL of nuclease-free water with the nCoV_IP2 primers (Forward 5'-ATG AGC TTA GTC CTG TTG-3'; Reverse 5'-CTC CCT TTG TTG TGT TGT-3').

**TCID50 quantification**

Culture supernatants were thawed and serial diluted (10-fold) from $10^{-1}$ to $10^{-8}$ in DMEM. Six replicates of viral dilutions (50 µL each) were seeded in flat bottom 96-well plates and mixed with 12,000 VeroE6 cells in DMEM-3% FBS (150 µL). After 4 days at 37°C, cells were fixed in 4% PFA for 30 min at room temperature (RT). PFA was then replaced by a crystal violet solution for 15 min at RT and rinsed with water. The TCID50 was determined as the lowest viral concentration inducing cell lysis (no crystal violet staining) in 100% of the 6 replicates.

**Transepithelial electical resistance (TEER) measurement**

The apical side of transwell cultures was washed for 20 min at 37°C in Mucilair$^{TM}$ medium. Transwell were then transferred in a new 24-well plate and DMEM medium was added to both the apical (200 µL) and basal (700 µL) sides. The TEER was then measured using an Evom3 ohmmeter (World Precision Instruments).

**Epithelium permeability assay**

The apical side of transwell cultures was washed for 20 min at 37°C in Mucilair$^{TM}$ medium, and transwell were then transferred in a new 24-well plate. Samples were then quickly rinsed with PBS on both apical and basal sides, and then and incubated in DMEM without phenol-red (Gibco). Dextran-FITC (4kDa, Sigma-Aldrich) was prepared at 1 mg/mL in DMEM without phenol-red, and 200 µL were added on the apical side. After 30 min at 37°C, the basal medium was harvested, and FITC fluorescence was measured using an EnSpire luminometer (Perkin Elmer). Medium alone and a Dextran-FITC solution (10 ng/mL) were used as negative and positive controls, respectively.

**LDH cytotoxicity assay**

Diluted culture supernatants (1:25) were pre-treated with Triton-X100 1% for 2 h at RT for viral inactivation. Lactate dehydrogenase (LDH) dosage was performed using the LDH-Glo™ Cytotoxicity Assay kit (Promega) following manufacturer's instructions. Luminescence was measured using an EnSpire luminometer (Perkin Elmer).

**Cytokine measurements**

Culture supernatants were pre-treated with Triton-X100 1% for 2 hr at RT for viral inactivation. IFN-β concentrations were quantified with a Simoa digital ELISA developed with Quanterix Homebrew kits as previously described (Rodero et al., 2017). Briefly, the 710322-9 IgG1-κ, mouse monoclonal antibody (PBL Assay Science) was used as a capture antibody to coat paramagnetic beads (0.3 mg/mL), the 710323-9 IgG1-κ mouse monoclonal antibody (PBL Assay Science) was biotinylated (biotin/antibody ratio = 40/1) and used as the detector antibody, and recombinant IFN-β protein (PBL Assay Science) was used as a reference to quantify IFN-β concentrations. The limit of detection (LOD) for IFN-β was 1.7 pg/mL. IFN-α2 was detected with a Simoa assay with a LOD of 2 fg/mL, as previously described (Hadjadj et al., 2020). Additional cytokines were measured with a commercial Legendplex bead-based immunoassay

(Biolegend), following manufacturer's instructions. Samples were analyzed on a LSRII flow cytometer (BD Biosciences).

**Immunofluorescence staining**

Reconstituted epithelia: MucilAir™ cultures were washed twice with PBS, fixed in 4% PFA for 30 min at RT, washed again twice with PBS and stored in PBS at 4°C until staining. Transwell membranes pieces were cut using a scalpel blade and staining steps were performed at RT on membrane pieces placed in 10 µL drops on parafilm.

For surface SARS-CoV-2 spike staining, membranes were blocked in PBS-0.1% Tween - 1% BSA - 10% FBS – 0.3 M glycine for 30 min, then incubated with mouse-anti-Spike antibody (kindly provided by N. Escriou, Institut Pasteur) at 1 µg/mL in PBS-0.1% Tween - 1% BSA for 30 min, followed by 3 washes of 5 min in PBS. Secondary anti-mouse antibody conjugated to AF488 (A-11001; Invitrogen) or AF555 (A-21422; Invitrogen) was added at 1:400 in PBS-0.1%Tween -1%BSA for 30 min, followed by 3 washes of 5 min in PBS and fixation in 4% PFA for 30 min at RT.

For intracellular staining, samples were first permeabilized with PBS-0.5% Triton for 20 min at RT and then blocked in PBS-0.1% Tween - 1% BSA - 10% FBS - 0.3 M glycine for 30 min. Samples were incubated with conjugated primary antibodies diluted in PBS -0.1% Tween - 1% BSA for 1 h at RT or overnight at 4°C, followed by 3 washes of 5 min in PBS. Samples were counterstained with Hoechst, followed by 3 washes of 5 min in PBS, and mounted in Fluoromount-G (Invitrogen) before observation with a STELLARIS (Leica Microsystems) or LM710 (Zeiss) confocal microscope.

Primary antibodies used were: rabbit anti-β-IV-tubulin-AF488 (ab204003; Abcam), rabbit anti-β-IV-tubulin-AF647 (ab204034; Abcam), rabbit anti-cytokeratin-5-AF647 (ab193895; Abcam), rabbit anti-mucin 5AC-AF555 (ab218714; Abcam), and rabbit anti-ZO-1(40-2200; Invitrogen).

Hamster tissues: Tracheas were aseptically harvested and fixed one week in 4% PFA, then washed in PBS and dehydrated in 30% sucrose. They were then embedded in O.C.T compound (Tissue-Tek), frozen on dry ice, and cryostat-sectioned into 20 µm-thick sections. Sections were rinsed in PBS, and epitope retrieval was performed by incubating sections for 20 min in citrate buffer (pH 6.0) at 96°C for 20 min. Sections were blocked in PBS-10% goat serum - 4% FBS - 0.4% Triton for 2 h at RT, then incubated with conjugated primary antibodies rabbit anti-$\beta$-IV-tubulin AF488 (ab204003; Abcam) and rabbit anti-cytokeratin-5 AF647 (ab193895; Abcam) at 1:100 in PBS-4% goat serum - 4% FBS - 0.4% Triton overnight at 4° C.

Sections were then counterstained with Hoechst, rinsed thoroughly in PBS, and mounted in Fluoromount-G (Invitrogen) before observation with a Zeiss LM 710 inverted confocal microscope.
Immunostaining was also done on whole pieces of hamster tracheas. Small samples (4 mm x4 mm) of tracheal tissue fixed in 4% PFA were microdissected, washed three times in PBS and blocked in PBS supplemented with 20% normal goat serum and 0.03% triton X-10 for 1 h at RT, then incubated overnight at 4° C with primary antibodies: mouse anti-Spike (1:100, provided by N. Escriou, Institut Pasteur) and rabbit conjugated-anti- $\beta$ -IV-tubulin-AF647 (ab204034; Abcam) in PBS supplemented with 2% BSA. The samples were rinsed three times in PBS, and incubated for 1 h at RT with secondary antibody ATTO-488-conjugated goat anti-mouse (1:500 dilution, in PBS - 2% BSA; Sigma-Aldrich). Sample where then mounted in Fluorsave (Calbiochem) and observed with a Zeiss LSM 710 confocal microscope.

**Confocal laser scanning microscopy**

MucilAir™ stained samples were visualized primarily with a STELLARIS 8 inverted microscope (Leica Microsystems, Mannheim, Germany). A super continuum white light laser tunable between 440 and 790nm was used for excitation and focused through an HC PL APO CS2 40x NA 1.1 water immersion or a HC PL APO CS2 63x NA 1,3 glycerol immersion objective. Emission signals were captured with Power HyD Detectors. The system was controlled with Leica Application Suite X (LAS) software. 3D images were directly processed using LAS X 3D module. The excitation wavelength and detection window are given in parentheses for each fluorophore: Hoeschst (405 nm; 410-470 nm); AlexaFluor488 (499 nm; 504-533 nm); AlexaFluor555 (553 nm; 558-600nm); AlexaFluor647 (653 nm; 658-700 nm).
Samples of hamster trachea were imaged with an inverted Zeiss LSM 710 confocal microscope. Z-stack images of tissue sections and whole mount samples were acquired with a Plan Apochromat 20x/0.8 Ph2 M27 lens or a Plan Apochromat 63x/1.4 N.A. oil immersion lens (Carl Zeiss).

**Scanning electron microscopy**

Samples were fixed with 2.5% glutaraldehyde in 0.1 M cacodylate buffer for 1h at RT and then for 12 h (MucilAir™ samples) or one week (hamster trachea samples) at 4°C to inactivate SARS-CoV-2. Samples were then washed in 0.1 M cacodylate buffer and several times in water, and processed by alternating incubations in 1% osmium tetroxide and 0.1 M thiocarbohydrazide (OTOTO), as previously described (Furness et al., 2008). After dehydration by incubation in increasing concentrations of ethanol, samples were critical point dried, mounted on a stub, and analyzed by field emission scanning electron microscopy with a Jeol JSM6700F microscope operating at 3kV.

**Transmission electron microscopy**

Samples were fixed with 2.5% glutaraldehyde in DMEM medium with 10% fetal calf serum for 1h at RT, and then overnight at 4°C. Fixed samples were washed three times for 5 min in PBS buffer, post-fixed for 1 h in 1% osmium tetroxyde solution, and rinsed with distilled water. Samples were dehydrated through a graded series of ethanol solutions (25, 50, 75, 95 and 100%) and embedded in epoxy resin. Samples were sectioned with an UC7 Leica ultramicrotome (Leica, Wetzlar, Germany) at a thickness of 70 nm, and transferred on 200 Mesh Square Copper grids coated with Formvar and carbon (FCF-200-Cu50, Delta Microscopy). Samples were then stained with 4% uranyl acetate and counterstained with lead citrate. Images were recorded with a TECNAI SPIRIT 120kV microscope with a bottom-mounted EAGLE 4K x 4K Camera, (ThermoFisher Scientific, Waltham, MA, USA).

**Image analysis**

Quantitative image analysis of the ZO-1-associated tight junction pattern

The surface of the epithelium contained in each microscopy z-stack was extracted using the Zellige program (C. Trébeau, R. Etournay, manuscript in preparation): the tissue 3D-manifold is determined based on both the detection of intensity maxima along the z direction and the spatial correlation of adjacent maxima along the x, y directions, with a maximum z difference $\Delta z=\pm1$ between adjacent maxima. The 3D manifold is then projected onto a plane. The TissueMiner software was then used to segment and quantify the cell apical surface area and the cell neighbor number (Etournay et al., 2016). The Wilcoxon test was used to compare the median cell area between conditions. The Welch corrected t-test was used to compare the average neighbor number between conditions.

Quantifications of β-tubulin+ area in epithelial cultures

Confocal images of samples stained with anti-β-IV-tubulin were analyzed. Images were visualized by Z-stack projection followed by thresholding (0.06) of the tubulin signal. The tubulin+ area was measured using the FIJI software, with the area measurement tool. The area was normalized to the total area of the picture.

Quantification of β-tubulin depth intensity profile in cilia

Samples were imaged at 63x magnification using the STELLARIS 8 microscope (Leica Microsystems).

Cell segmentation: An ImageJ user-guided tool was developed to extract single cells in the epithelium. Three cell categories were defined: cell from a mock-infected sample (mock), cell expressing the spike antigen in an infected sample (spike+), and bystander cell not expressing the spike antigen in an infected sample (spike-). A dataset of 79 mock, 85 spike+, and 71 spike- cells was extracted from 2 mock samples and 4 SARS-CoV-2-infected samples.

Averaged cell intensity profiles: For each cropped cell, the intensities of β-tubulin and spike were averaged along the depth axis. The resulting intensity profiles for all cells were then re-aligned using as a reference the proximal intensity peak of β-tubulin located just below the plasma membrane. The average intensity profiles of β-tubulin and spike along the cell depth axis was then computed for each of the 3 cell categories.

β-tubulin distal to proximal peak intensity ratio: The averaged β-tubulin profiles showed two peaks, with a distal peak corresponding to cilia and a proximal peak located just below the plasma membrane, which was used for realignment. The proximal peak corresponded to the area where basal bodies anchor cilia into the cytoplasm. The ratio between the intensities of the distal peak and the proximal peak was computed for each cell.

Colocalization analysis: Images were acquired at the Z level where the spike signal was maximal, corresponding to the area just above the plasma membrane. Images were deconvoluted with Huygens Professional version 19.04 (Scientific Volume Imaging, The Netherlands, http://svi.nl). Thresholded Mander's colocalization coefficients between the β-tubulin and spike markers were then computed using Otsu algorithm (Otsu, 1979).

<u>Quantification of basal cell height</u>

Tissue level correction: The reconstituted epithelia were grown on inserts that were not perfectly flat, due to experimental constraints. It was necessary to correct for local insert deformation before measuring the height of basal cells within the epithelia. Due to the insert strong autofluorescence in both the spike and cytokeratin-5 channels, insert elevation could be measured by finding the depth value of the maximal intensity of both signals multiplied, defined as $Z_{insert}(x,y) = argmax_z(I_{spike}(x,y) \times I_{btub}(x,y))$. This value provided the insert elevation map used to correct for tissue deformation. The map was blurred with a gaussian filter of sigma=3 to smooth the results and remove possible small noise.

Cytokeratin-5 density profile: The cytokeratin-5 pixel density profile was computed by first smoothing the signal using a sigma=1.5 gaussian blur and then thresholding the signal using Otsu algorithm (Otsu, 1979). The Cytokeratin-5 density profile for each sample was computed along the Z-axis by measuring the ratio of thresholded pixels over total pixels per plan. The profiles were realigned using the 2/3 of maximum intensity and averaged together. The significance of the difference between the profiles obtained from mock and infected samples was evaluated with a Welch t-test comparing profile area under the curve, using the trapezoidal numerical integration algorithm.

Quantification of ciliated area in SEM images

Ciliated areas in SEM images were selected and masked in the Adobe Photoshop v21.1.3. software, using the object selection tool, which uses machine learning to automatically shrink wrap an object. Selection of each ciliated area was refined manually with the lasso tool. The masked surface was then measured in the Fiji software (Schindelin et al., 2012).

**Mucociliary clearance**

Experiments were performed at 7 dpi to allow mucus layer reconstitution after infection. 10 µL of 30 µm polystyrene beads (Sigma Aldrich) diluted 1:3 in PBS were added to the apical side of transwell cultures. Bead movements were recorded at 37°C in the brightfield channel of a Biostation inverted microscope (Nikon), taking 1 picture every 2 s during 1 min. A minimum of 7 fields per sample were recorded within the first 15 min after addition of beads.

For image analysis, beads were first manually dot-marked using pencil tool from the FIJI software. Tracks were generated and analyzed using the TrackMate FIJI plugin v5.2.0 (Tinevez et al., 2017). Dot-marked beads were automatically detected with LoG detector (estimated diameter = 20 µm; threshold = 5 to 8). Tracks were then generated using the Simple LAP tracker (gap closing max frame gap = 0), adjusting linking max distance (50 to 200 µm) and gap-closing max distance (15 to 100 µm) for each field depending on speed and bead density. The resulting tracks statistics included speed, XY movements, and track straightness parameters. Tracks composed of 3 dots of less were removed from the analysis. Examples of track images for Figure 5 were generated with the Imaris software (Oxford Instruments).

**3- STATISTICAL ANALYSIS**

Statistical analyses were performed with the Prism software v8 (GraphPad). The non-parametric Mann-Whitney test was used, except when data was estimated to be normally distributed (Fig. 3B, Fig. 5E-F, and Fig. 6C). Statistical significance was assigned when p values were $< 0.05$. Box-whisker plots show median (horizontal line), interquartile range (box), and 1.5 times the interquartile range (whiskers). The nature of statistical tests and the number of experiments or animals (n) are reported in the figure legends.

_______

**LEGENDS TO THE SUPPLEMENTAL FIGURES**

**Figure S1: Quantitative image analysis of the ZO-1-associated tight junction pattern**

Reconstituted epithelia were labeled with a ZO-1 antibody (red) to detect the tight junction pattern. Confocal images were analyzed with a dedicated software pipeline to quantify cell shape and cell neighbor numbers.

**(A-H)** Examples of ZO-1 image analyzes for mock (A-D) and SARS-CoV-2 infected (E-H) epithelia at 4 dpi. Original microscopy z-stack are visualized in (A, E). The surface of the epithelium is extracted and projected onto the (x,y) plane using the Zellige program (C. Trébeau, R. Etournay, in preparation) (B,F). Cell segmentation and subsequent quantifications are performed with the TissueMiner software using the data processed with Zellige. The cell area size (C, G) is color-coded in the range of 0-100 $\mu m^2$. The cell neighbor number pattern (D, H) is color-coded according to the number of neighbors (polygon class). Scale bars measure 20 µm.

**(I)** Comparison of cell area distribution in mock and infected epithelia at 2, 4, and 7 dpi. The Wilcoxon test was used to compare the median cell area between conditions.

**(J)** Comparison of the distribution of cell neighbor numbers in mock and infected epithelia at 2, 4, and 7 dpi. The Welch corrected t-test was used to compare the average neighbor number between conditions (P values reported next to vertical arrows).

**Figure S2: SARS-CoV-2 induces a cytopathic effect in reconstituted human bronchial epithelia**

**(A)** Quantification of the lactate dehydrogenase enzyme (LDH) in apical supernatants to evaluate cell death in mock and SARS-CoV-2 infected epithelia (mean ± SD for n=2 independent experiments).

**(B)** 3D confocal image of a SARS-CoV-2 infected epithelium showing nucleus extrusions (Hoechst, blue) above the layer of ciliated cells (β-tubulin+, red) in an area with viral production (spike+, green).

**(C)** Scanning electron microscopy (SEM) image of a rounded extruded cell (left). An enlargement shows the presence of viral particles (vp) at the cell membrane (right).

**Figure S3: SARS-CoV-2 occasionally infects basal and secretory cells**

**(A)** Confocal image of a rare infected basal cell (cytokeratin 5+ and spike+; arrowhead) in a reconstituted epithelium at 2 dpi.

**(B)** Goblet cells (MUC5AC+, cyan) in a reconstituted epithelium at 2 dpi do not express the spike antigen (green).

**(C**) SEM image of a secretory cell with characteristic membrane pores and abundant viral particles (vp) at its surface. An enlargement of the framed area is shown to the right.

**(D)** SEM image of a transitional cell showing both secretory vesicles (sv) and cilia (ci). The enlargement to the right shows the presence of viral particles (vp) at the cell surface.

**Figure S4: Image analysis of cilia**

**(A)** Example confocal image showing the distribution of β-tubulin IV (red) and SARS-CoV-2 spike (green) along the depth axis of a reconstituted epithelium at 2 dpi.

**(B, C)** Analysis of spike and β-tubulin colocalization in infected cells at 2 dpi.

(B) shows a representative infected cell viewed in the x,y plane at membrane level. Little overlap is seen between cilia (red spots) and spike+ areas (green spots), as shown by the limited extent of the yellow area. A thresholded image (bottom right) was used to compute colocalization coefficients.

(C) Measurement of Mander colocalization coefficient for n>70 infected cells. The Mander Tubulin coefficient measures the fraction of β-tubulin+ intensity that is located in spike+ pixels. Conversely, the Mander Spike coefficient measures the fraction of spike intensity that is located in β-tubulin+ pixels. Both coefficients are low, indicating minimal colocalization of the two markers.

**(D)** TEM image of cilia seen in transverse section, in an uninfected epithelium. In each cilium, 9 peripheral microtubule doublets and a central microtubule pair are visible.

**(E)** TEM image of an infected epithelium at 4 dpi showing mislocalized basal bodies (bb) close to or engulfed in vacuoles, viral particle (vp) accumulation in intracellular vesicles, and an electron-dense inclusion body (ib).

**Figure S5: Image analysis of basal cells**

**(A)** Image correction for local height deformation. The tissue sample was grown on an insert that was not entirely flat due to experimental constraints (left panel, top). The height of basal cells (cytokeratin-5+, yellow) is skewed by insert deformation, as shown in an orthogonal projection (left panel, bottom). The insert is visible on confocal images due to autofluorescence, so that an insert deformation map can be generated (middle panel). The map is then used to correct images for tissue deformation (right panel).

**(B)** SEM images of basal cells in mock (left) and infected (right) epithelia at 7 dpi. Two representative images are shown for each condition.

**Figure S6: SARS-CoV-2 infection targets ciliated cells in the trachea of infected Syrian hamsters**

Confocal images of the tracheal epithelium of mock infected (left) or SARS-CoV-2 infected (middle and right) hamsters at 4 dpi. Labeling for the SARS-CoV-2 spike (A, green), the ciliated cell marker β-tubulin IV (B, red), or both markers (C) are shown. The tracheal epithelium obtained in the infected animal show various degrees of cilia destruction (middle, right) depending on the area.

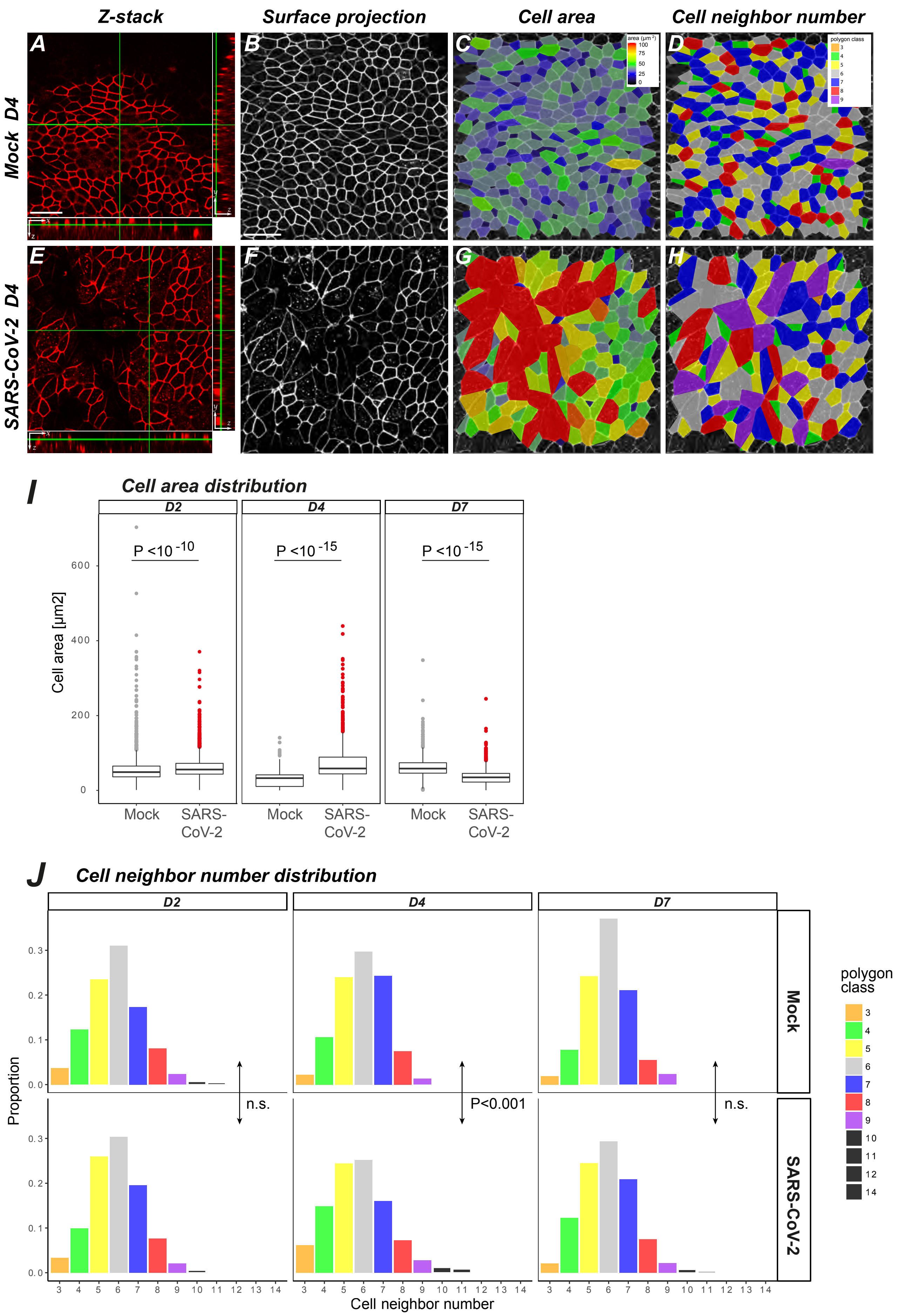
Z-stack
Surface projection
Cell area
Cell neighbor number
Mock D4
SARS-CoV-2 D4
A
B
C
D
E
F
G
H
I
Cell area distribution
D2
D4
D7
P <10 -10
P <10 -15
P <10 -15
Cell area [µm2]
0
200
400
600
Mock
SARS-CoV-2
J
Cell neighbor number distribution
Proportion
0.0
0.1
0.2
0.3
n.s.
P<0.001
n.s.
Mock
SARS-CoV-2
Cell neighbor number
3 4 5 6 7 8 9 10 11 12 13 14
polygon class
14

*Figure S1*

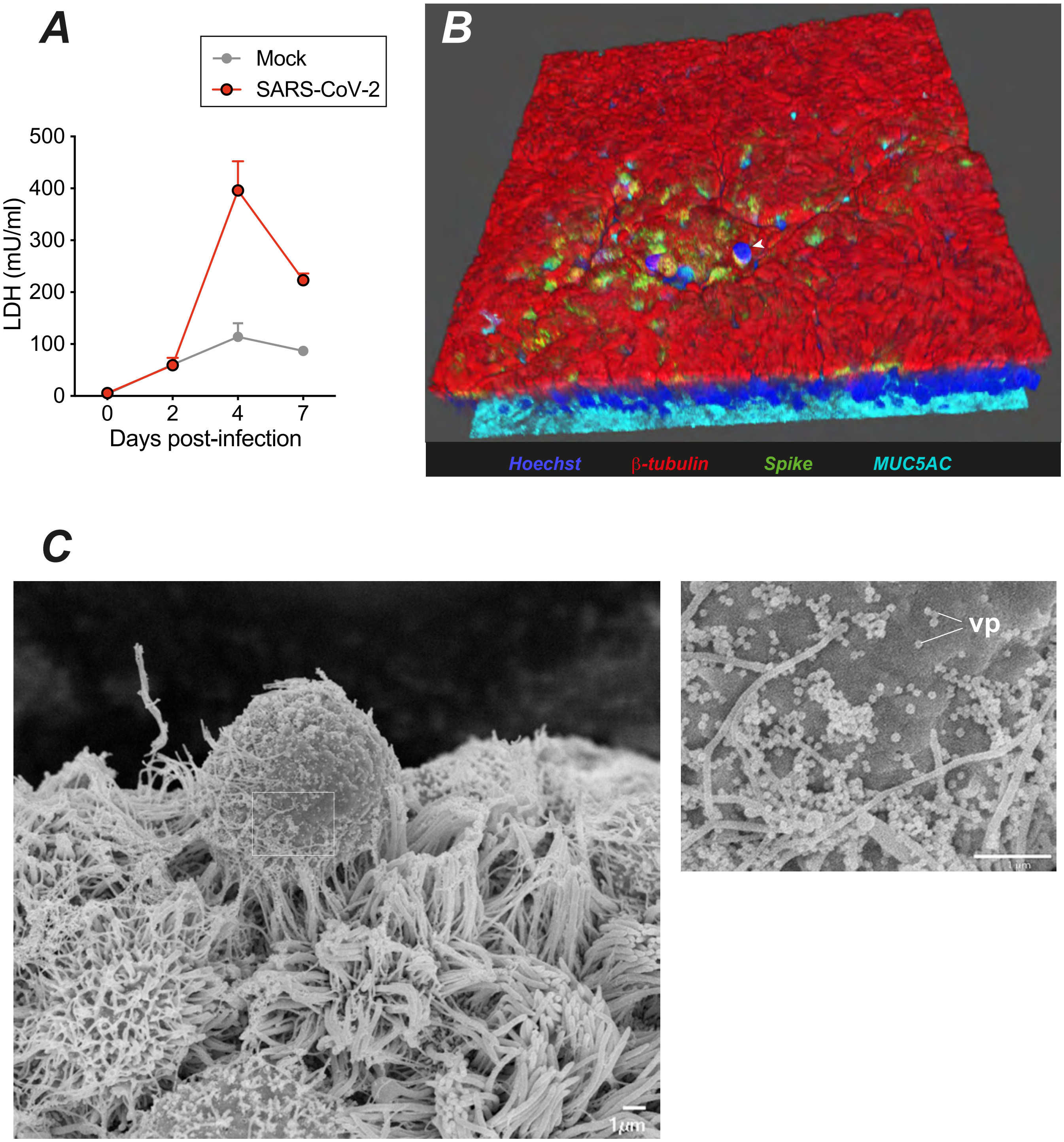
A
Mock
SARS-CoV-2
500
400
300
200
100
0
LDH (mU/ml)
0
2
4
7
Days post-infection
B
Hoechst
β-tubulin
Spike
MUC5AC
C
vp
1µm

*Figure S2*

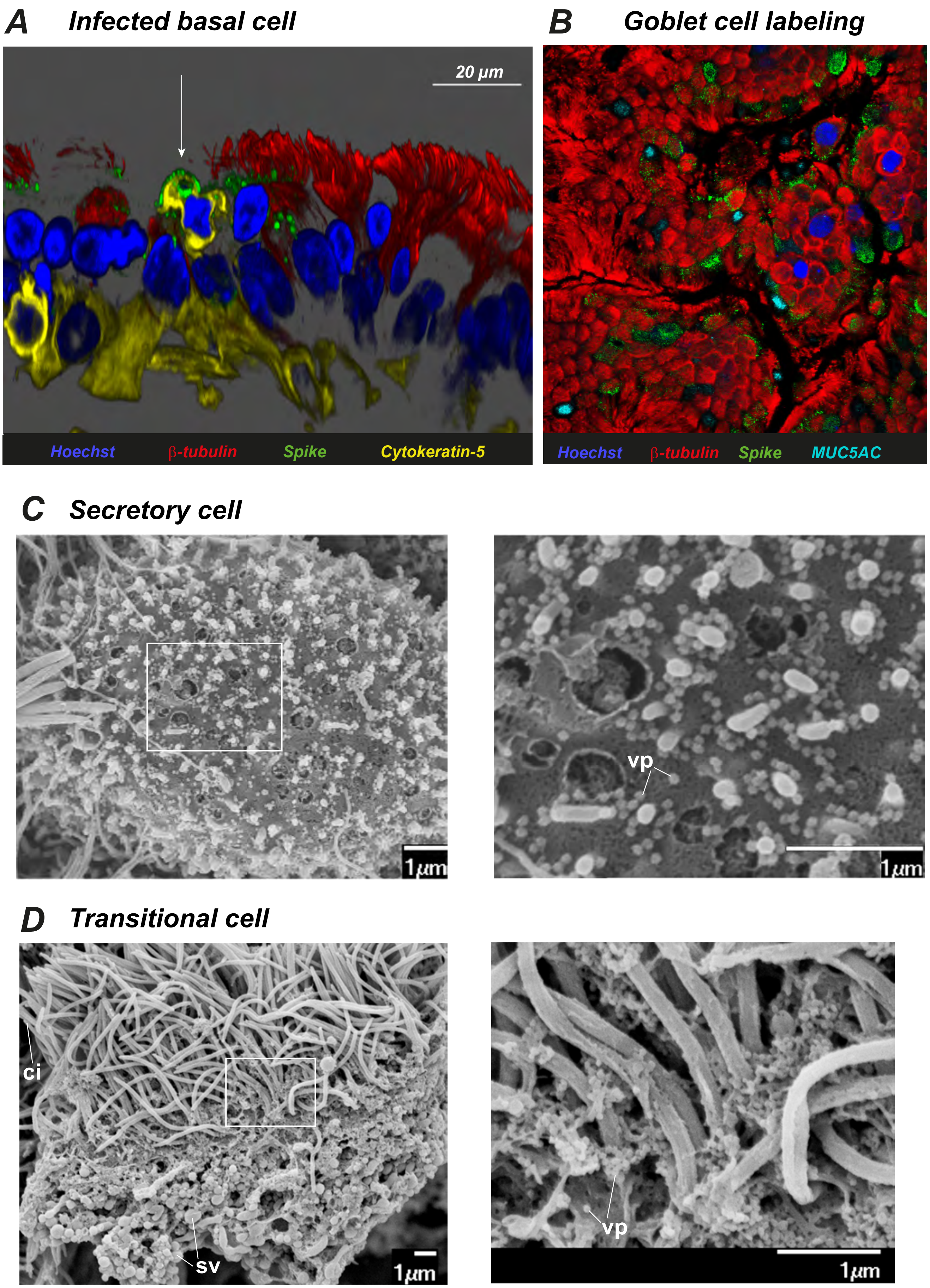
A
Infected basal cell
20 µm
Hoechst
β-tubulin
Spike
Cytokeratin-5
B
Goblet cell labeling
Hoechst
β-tubulin
Spike
MUC5AC
C
Secretory cell
1µm
vp
1µm
D
Transitional cell
ci
sv
1µm
vp
1µm

Figure S3

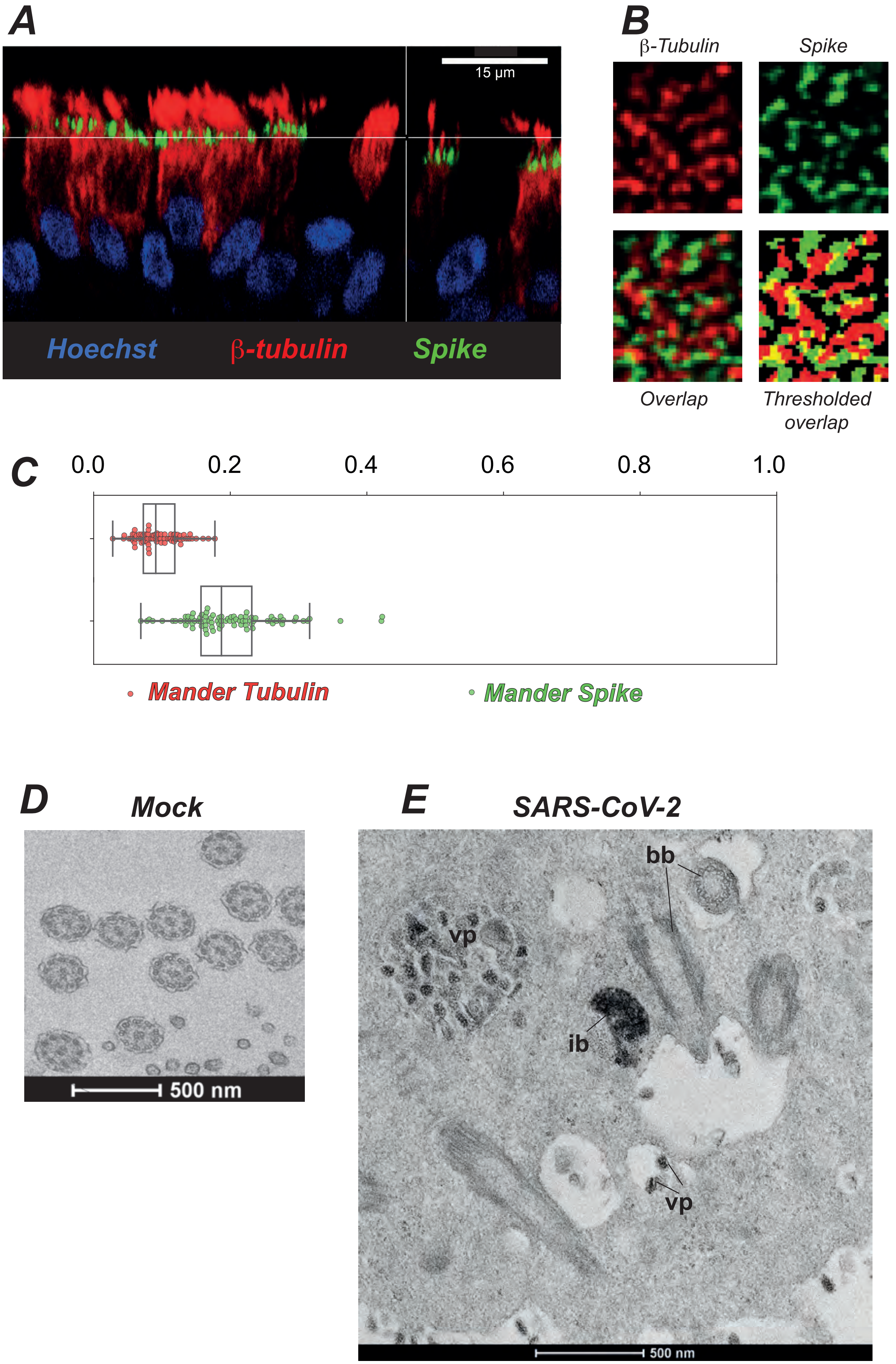
A
15 µm
Hoechst
β-tubulin
Spike
B
β-Tubulin
Spike
Overlap
Thresholded overlap
C
0.0
0.2
0.4
0.6
0.8
1.0
Mander Tubulin
Mander Spike
D
Mock
500 nm
E
SARS-CoV-2
bb
vp
ib
vp
500 nm

Figure S4

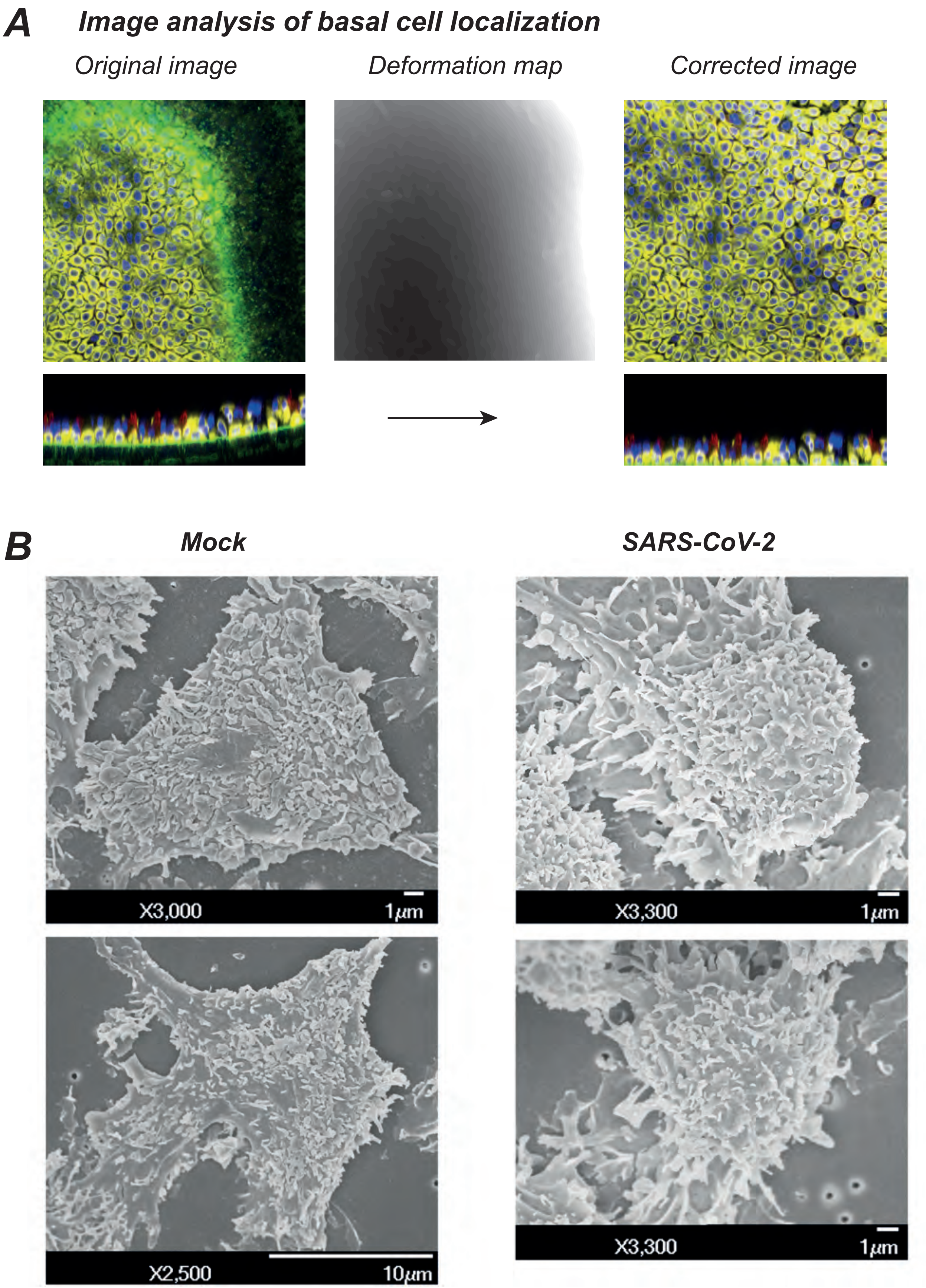
A
Image analysis of basal cell localization
Original image
Deformation map
Corrected image
B
Mock
SARS-CoV-2
X3,000
1μm
X3,300
1μm
X2,500
10μm
X3,300
1μm

Figure S5

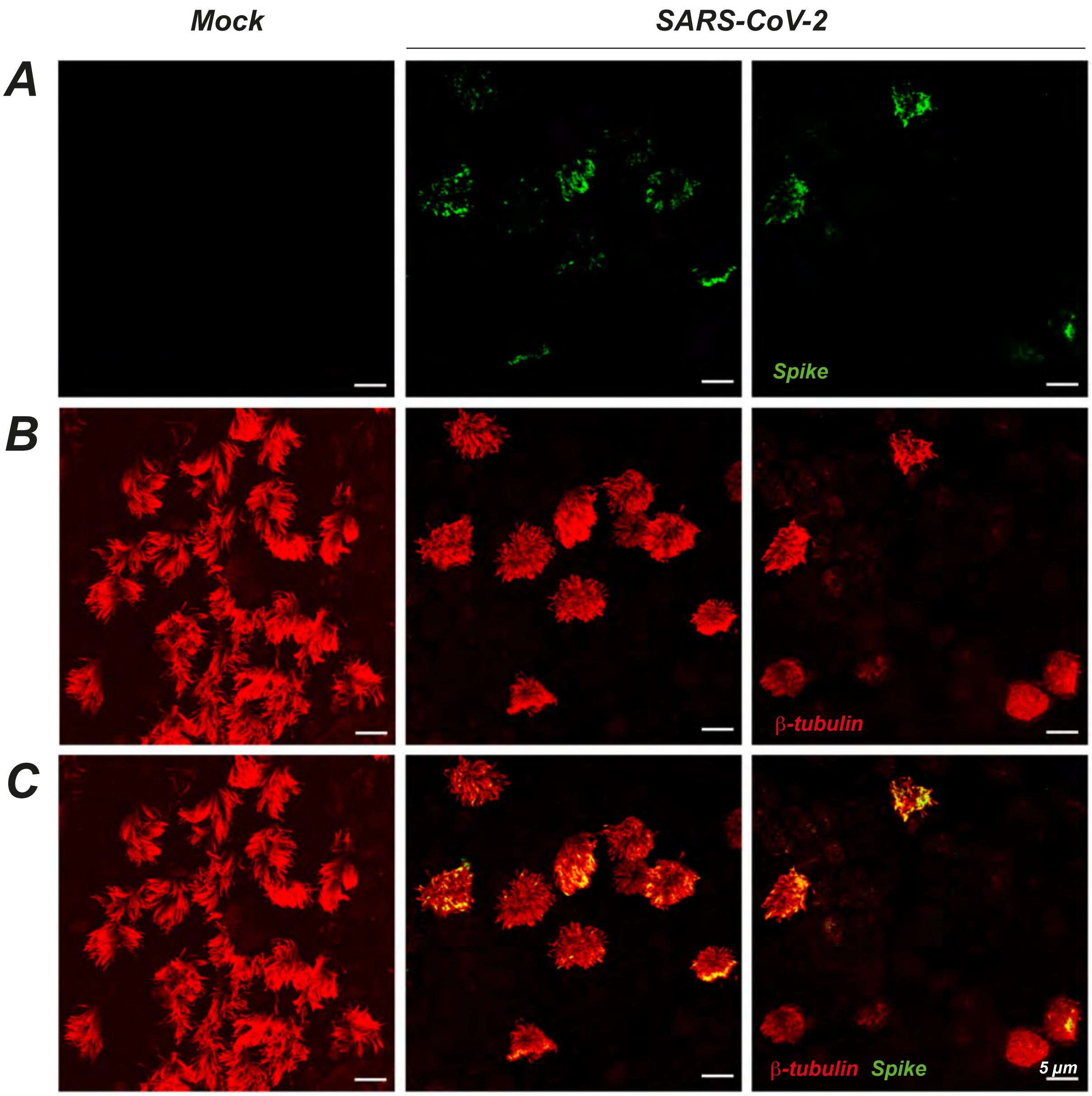

Figure S6
